# Dyslipidemic SPTLC3 Integrates Bile Acid-FXR Signaling with Sphingolipid Remodeling in MASLD

**DOI:** 10.1101/2025.08.25.672208

**Authors:** Michele Visentin, Zhibo Gai, Anna Kovilakath, Francesca De Luca, Grigorios Panteloglou, Alaa Othman, Mingxia Liu, Francesca Barone, Imme Garrelfs, Joanne Verheij, Belle V. van Rosmalen, Ting Gui, Bin Yan, Zhiyong Zhang, Andreas J. Hülsmeier, Sophia L. Samodelov, Essa M. Saied, Mary Gonzalez Melo, Arnold von Eckardstein, Christoph Arenz, John S. Parks, L. Ashley Cowart, Wolfgang Thasler, Gerd A. Kullak-Ublick, Thorsten Hornemann, Museer A. Lone

## Abstract

Molecular mechanisms driving metabolic disease pathogenesis remain poorly understood. Genetic and functional studies implicate SPTLC3 with dyslipidemia. SPTLC3 synthesizes atypical long-chain bases (LCB) precursors for sphingolipid production. We demonstrate significant *SPTLC3* expression in human liver. ORMDL1-3 regulate SPTLC3 post-translationally in hepatic and non-hepatic cells. Independently, farnesoid X receptor (FXR) represses hepatic *SPTLC3* transcription via a negative promoter element. In mice, high-fat diet (HFD) induced whereas bile acids normalized hepatic *SPTLC3* transcription. SPTLC3 derived LCBs in plasma originate from the liver and are elevated in HFD-fed mice and in patients with metabolic dysfunction-associated steatotic liver disease (MASLD). Cross-species comparison revealed marked differences in LCB composition between mice and humans. Notably, omega-3-methylsphingosine (meC_18_SO) was significantly associated with MASLD in humans but undetectable in mice. In Huh7 cells, meC_18_SO enhanced complex II and IV activity, oxygen consumption, and mitochondrial ROS content. FXR-SPTLC3 axis and presented findings have potential implications for future translational research.

## Introduction

Metabolic dysfunction-associated steatotic liver disease (MASLD), a recently adopted term replacing non-alcoholic fatty liver disease (NAFLD)[1], is a progressive disorder characterized by aberrant lipid accumulation in the liver that can lead metabolic dysfunction-associated steatohepatitis (MASH), fibrosis, cirrhosis, and hepatocellular carcinoma. MASLD is a major global health burden and an independent risk factor for type 2 diabetes mellitus (T2DM) and coronary heart disease [2]. Despite extensive investigation, the molecular mechanisms underlying MASLD or its link to the systemic metabolic dysfunction remain poorly defined.

Sphingolipids (SLs) are increasingly recognized as important contributors to metabolic disorders, including MASLD [3, 4]. SL are essential structural and signaling lipids, composed of an identifying aliphatic amino alcoholic long-chain base (LCB), conjugated to a fatty acid and variable functional head groups [5]. LCB formation, from acyl-CoA and L-serine by serine-palmitoyltransferase (SPT), is the first step in the de novo SL synthesis. Mammalian genome encodes three SPT subunits, namely *SPTLC1, SPTLC2,* and *SPTLC3* (SPTLC isoforms). The minimal SPT active enzyme may consist of SPTLC1 and SPTLC2 or SPTLC3 [6]. Ubiquitously expressed SPTLC1 and 2 display the highest affinity for palmitoyl-CoA and L-serine, generating C_18_ LCB precursors for canonical SL (Supplementary Fig. S1A). In contrast, SPTLC3 shows variable tissue expression and generates LCBs of variable chain lengths, including the branched chain, omega-3-methylsphingosine (meC_18_SO) in humans (Supplementary Fig. S1B) [6, 7].

SPT is also the rate-limiting step in *de novo* SL synthesis. Post-translational regulation of the SPTLC1/SPTLC2 complex is mediated by ceramide induced feedback inhibition, involving ORMDL1-3 proteins (ORMDLs) [8, 9]. Structural [10, 11] and mechanistic [12–16] details regarding the ORMDL mediated regulation of homologous SPTLC1/2 complexes across species have emerged in recent years. Perturbation of SPT-ORMDL interaction has severe pathophysiological consequences in humans. Mutations in SPTLC1 and SPTLC2 that impede SPT-ORMDL interaction and feedback regulation of the SL synthesis are associated with juvenile amyotrophic lateral sclerosis (jALS) [17–22]. In addition, ORMDLs appear to be implicated in the development of metabolic diseases [23]. ORMDL3-deficient mice exhibit enhanced weight gain, insulin resistance, and impaired adipose thermogenesis [24]. However, the mechanisms through which ORMDLs drive metabolic dysfunction remain unclear.

Epidemiological and experimental studies increasingly implicate SPTLC3 in metabolic disease. *SPTLC3* genetic locus has been linked to dyslipidemia, hepatic and cardiovascular dysfunction [25–31]. In mouse models of MASLD, hepatic *SPTLC3* expression is elevated [32–34]. In humans and mice, *SPTLC3* expression correlates with disease progression to hepatocellular carcinoma in mice and humans [35, 36]. Whereas *SPTLC3* variants are associated with elevated LDL-Cholesterol [37, 38], SPTLC3 derived SL are present in high- and low-density lipoproteins (HDL and LDL) [6]. Hepatic expression of *SPTLC3* and genes that regulate lipoprotein particle secretion from liver are positively correlated [39]. Moreover, SPTLC3 likely contributes to *trans*-fat induced cardiovascular disease (CVD) [39]. SPTLC3 is directly associated with cardiac diseases [40]. SPTLC3 transcription is upregulated, under the influence of HIF1, in ischemic cardiomyopathy [40]. Alternately, reduction in SPTLC3 levels counters oxidative stress, fibrosis, and hypertrophy associated with chronic ischemia in mice [40]. Together, these findings position SPTLC3 as a potential mechanistic link between hepatic dysfunction, dysregulated lipid metabolism, and CVD [41].

The bile-activated nuclear receptor, Farnesoid X receptor alpha (FXR, encoded by NR1H4) is an important therapeutic target in metabolic diseases, including MASLD [42, 43]. FXR shows highest expression in the liver and intestines [44]. Bile acids, also synthesized and recycled along the enterohepatic axis from the common substrate cholesterol, are physiological FXR activators [45–47]. Activated FXR dimerizes with Retinoid X Receptor alpha (RXRα, NR2B1) and binds to FXR response elements (FXREs) in the target genes. FRXEs are largely represented by hexanucleotide inverted repeats -AGGTCA-separated by a single nucleotide (IR-1) [48]. Yet, activated FXR can also act independent of RXRα or bind at the FXRE half-sites as well as at the non-canonical FXRE sequence sites [49–51]. FXR also acts through other nuclear receptors that then bind specific DNA elements in their target genes [52, 53]. In any case, bile acid activated FXR ensures stimulation or suppression of target genes encoding pace-limiting transporters or enzymes in lipid and glucose metabolism [42, 54–56]. FXR, for example, regulates cholesterol metabolism through regulation of HDL uptake by SR-B1 [57] and repression of the cholesterol 7α-hydroxylase (CYP7A) [52, 53], the rate-limiting enzyme of bile acid synthesis. Recent data suggests that bile acid/FXR signaling may also intersect with SL metabolism [58]. However, a direct mechanistic link between FXR activity and the *de novo* sphingolipid biosynthesis pathway has not been established.

A deeper understanding of SPTLC3 function and the mechanisms regulating its expression may provide valuable insight into the pathophysiology of metabolic diseases. Here, we investigated the regulation of SPTLC3 by ORMDL proteins and bile acid–activated FXR. We show that SPTLC3-containing SPT complexes are modulated post-translationally by ORMDLs in hepatic and non-hepatic cells. Independently, FXR represses SPTLC3 transcription in hepatocytes and mouse liver. Additionally, SPTLC3-derived long-chain bases are elevated in both MASLD patients and high-fat diet (HFD)-fed mice, and cross-species comparison reveals striking differences in SL profiles, particularly the presence of omega-3-methylsphingosine (meC_18_SO) in humans. These findings define a regulatory axis linking FXR signaling to SL metabolism and establish SPTLC3 as a relevant node in metabolic disease.

## Results

### ORMDL regulation of SPTLC3 containing SPT complex

We sought to systematically probe factors contributing to the regulation of SPTLC3 activity. First, we looked at posttranslational regulation of SPTLC3 by manipulating ORMDL expression in HEK293 cells. Also, diverse and often isobaric species pose a challenge in SL identification and quantification. Therefore, for direct assessment of SPT function, extracted lipids were first hydrolyzed to release LCBs that were then quantified by liquid chromatography mass spectrometry (LC-MS) [6, 59]. Additionally, as metabolic readout in these assays, *de novo* LCB synthesis was probed with D_3_, ^15^N-L-serine supplementation [6].

HEK293 cells endogenously express SPTLC1 and SPTLC2 subunits and form C_18_- and C_20_-LCB[6] (Supplementary Fig. S1A and B). However, SPTLC3 expression induced, in addition to C_18_SO, *de novo* synthesis of atypical LCBs, C_16_SO(d16:1), C_17_SO(d17:1) and C_20_SO(d20:1) as well as the branched LCB, meC_18_SO(d19:1) (Supplementary Fig. S2A and B). Principal component analysis (PCA) of *de novo* synthesized LCBs in HEK293 cells showed that cells expressing SPTLC3 clustered separately from wild-type, SPTLC1-, and SPTLC2-expressing cells (Fig. 1A). Previous studies have shown that only the simultaneous knockdown (KD) or co-expression of all three ORMDL isoforms leads to significant upregulation or repression, respectively, of *de novo* LCB synthesis [12, 60–62]. Therefore, siRNA-mediated KD of ORMDL1-3 was performed simultaneously in wild-type and SPTLC3-expressing HEK293 cells to assess SPT activity. ORMDL KD induced an increase in the sphinganine and sphingosine forms of C_18_- and C_20_-LCB in HEK293 wild type and SPTLC3 expressing cells (Fig. 1B). The relative change for LCB levels in HEK293 wild-type cells appeared higher compared to SPTLC3 expressing cells as the latter already showed a significant increase in de novo LCB synthesis (Supplementary Fig. S2A). Similarly, ORMDL KD led to a significant increase in atypical LCBs in SPTLC3 expressing cells (Fig 1B). However, ORMDL KD caused a relatively higher increase in the synthesis of saturated LCBs, C_16_SA (d16:0), C_17_SA(d17:0), C_18_SA (d18:0), C_20_SA(d20:0) and meC_18_SA(d19:0) (Fig. 1B).

**Figure 1:**
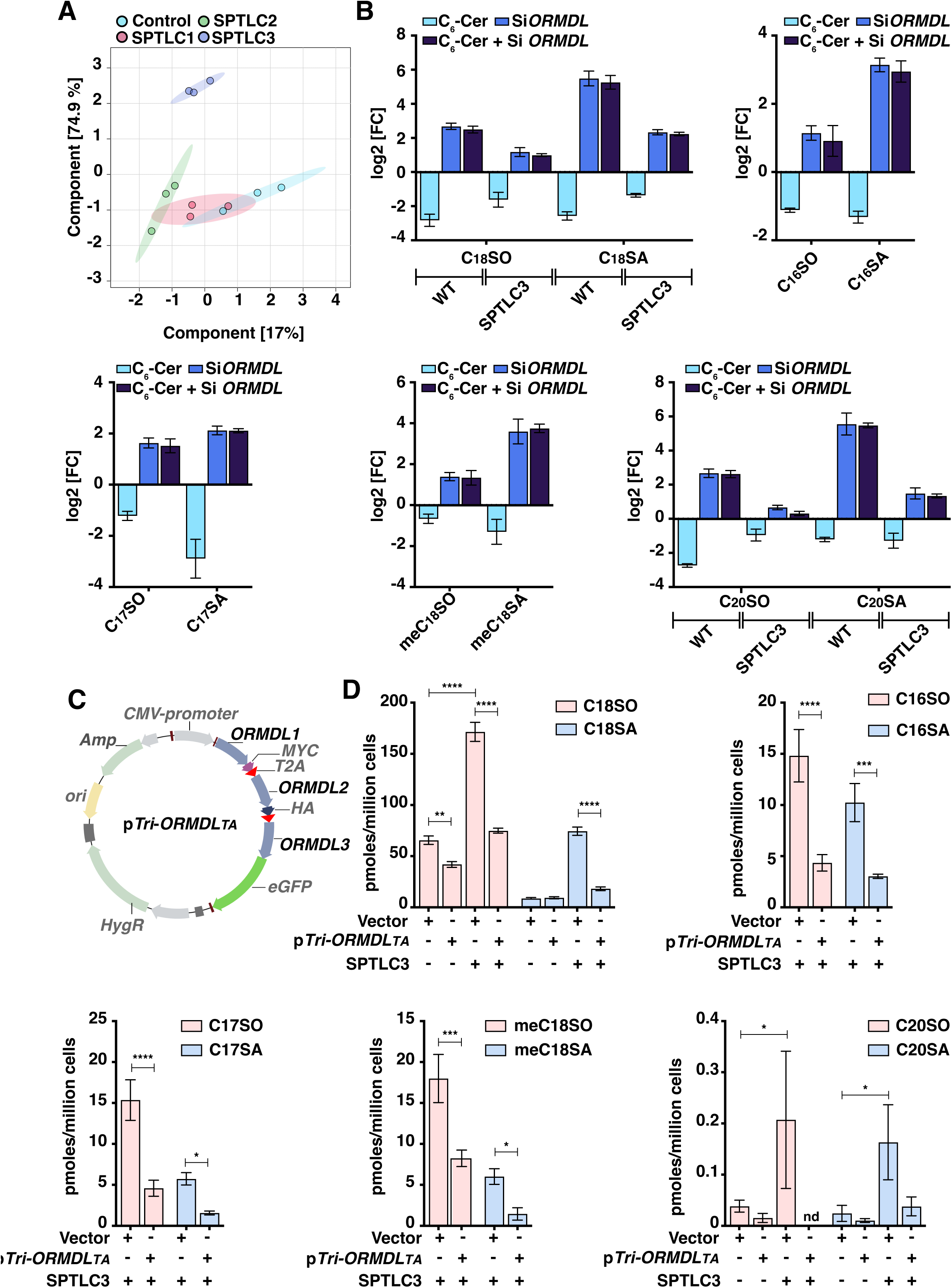
Post translational regulation of SPTLC3 by ORMDL1-3 proteins. (A) Principal component analysis (PCA) for long chain base (LCB) profiles in HEK293 over expressing SPTLC isoforms. (B) Levels of de novo produced C_18_, C_16_, C_17_, meC_18_, and C_20_ based LCB in wild type (WT) and SPTLC3 expressing HEK293 cells after ORMDL1-3 silencing, in the presence or absence of a ceramide analogue (C_6_-Cer). (C) Map depicting polycistronic ORMDL1-MYC, ORMDL2-HA and, ORMDL-GFP plasmid (pTri-ORMDL_TA_). (D) Levels of *de novo* produced LCB in vector or pTri-ORMDL_TA_ transfected wild type and SPTLC3 expressing HEK293 cells. PCA was performed with Metaboanalyst suite 5.0. Bars represent mean ± SD. Statistical significance was assessed by one-way ANOVA followed by Tukey’s correction (\**P* < 0.05, \*\**P* < 0.01, \*\*\**P* < 0.001, \*\*\*\**P* < 0.0001).

Further, the ceramide analogue, (C_6_-Cer), induces ORMDL mediated inhibition of the SPTLC1/SPTLC2 enzyme in HEK293 cells [18]. C_6_-Cer supplementation induced dose dependent inhibition of C_18_SO and atypical LCB synthesis in wild type as well as SPTLC3 expressing cells (Supplementary Fig. S2B). ORMDL KD abrogated C_6_-Cer mediated inhibition of LCB synthesis in both wild type and SPTLC3 expressing cells (Fig. 1B).

Next, we probed the effect of ORMDL over-expression on SPTLC3 activity. To achieve comparable expression, polycistronic *ORMDL1-3* coding sequences were expressed from a common promoter (Fig. 1C). To ensure translation of three distinct proteins from a common transcript, self-cleaving viral T2A peptides were inserted between the individual ORMDL coding sequences. This plasmid, designated p*Tri-ORMDL_TA_* (Fig. 1C) (Supplementary sequence file 1), was then transfected into HEK293 wild-type and SPTLC3-expressing cells. p*Tri-ORMDL_TA_* transfected cells showed a significantly decreased synthesis of C_18_SO(d18:1) and C_18_SA(d18:0) in both cell types, although relative to the vector transfected cells the decrease in SPTLC3 expressing cells appeared more pronounced (Fig. 1D). p*Tri-ORMDL_TA_* also induced significant reduction in the *de novo* synthesis of C_16_SO(d16:1), C_17_SO(d17:1), C_20_SO(d20:1) and meC_18_SO(d19:1) in SPTLC3 expressing cells (Fig. 1D).

Taken together, these data point to ORMDL-mediated posttranslational regulation of the SPTLC1-SPTLC3 enzyme complex.

### Bile acids regulate *SPTLC3* expression in hepatocytes

Given the association of SPTLC3 with hepatic dysfunction and transcriptional response to metabolic state [41], we wanted to examine its regulation in human hepatic cells. As SPTLC3 expression is tissue restricted, we wanted to check whether human liver cells express the enzyme. Single cell transcriptomic data from publicly available sources [63] showed significant *SPTLC3* expression in the major hepatic cell types, hepatocytes, hepatic stellate cells, Kupffer cells, and liver sinusoidal endothelial cells (Supplementary Fig. S3A). Interestingly, in hepatocytes, which represent around 70% of the hepatic cell population, *SPTLC3* expression exceeds that of *SPTLC2* (Supplementary Fig. S3A). For *in vitro* studies we chose human hepatoma-derived Huh7 cells. Analysis of our published RNA-Seq data [64] and qRT-PCR confirmed relative expression of SPTLC isoforms in Huh7 cells (Supplementary Fig. 3B and Fig. 2A). Similar analysis confirmed expression of all three ORMDL genes in hepatocytes and Huh7 cells (Supplementary Fig. 3A-B and Fig. 2A). Huh7 cells also synthesize SPTLC3 derived LCB, including the meC_18_SO [6]. Therefore, Huh7 cells represent a suitable cellular model for studying post-translational regulation of SPTLC3 in hepatocytes. As expected, concomitant siRNA-mediated KD of ORMDLs in Huh7 cells increased LCBs of all chain lengths (Fig. 2A and B). Consistent with the pattern seen in SPTLC3-expressing HEK293 cells, ORMDL knockdown in Huh7 cells led to a greater relative increase in saturated LCB species compared to mono-unsaturated LCBs.

**Figure 2:**
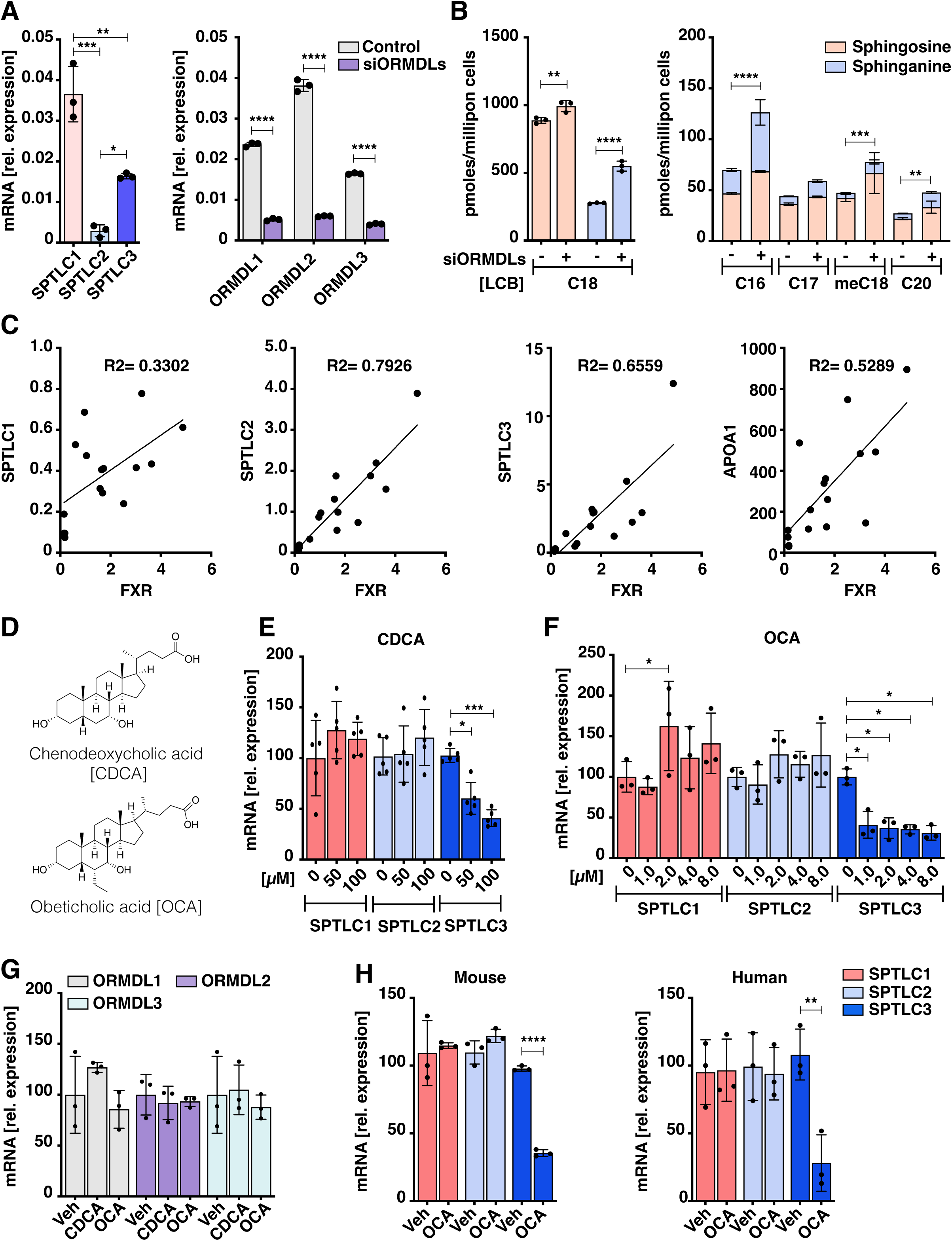
Bile acids regulate SPTLC3 expression in hepatocytes. (A) Relative SPTLC1-3 and ORMDL1-3 transcript levels in Huh7 cells. (B) Long chain base (LCB) levels in Huh7 cells after ORMDL silencing. (C) Correlation of SPTLC isoform and APOA1 expression with farnesoid X receptor (FXR) mRNA in liver tissues. (D) Structures of natural and semi-synthetic FXR agonists, chenodeoxycholic acid (CDCA) and obeticholic acid (OCA). (E-F) Relative SPTLC mRNA levels in Huh7 cells supplemented with CDCA or OCA. (G) Relative ORMDL mRNA level in Huh7 cells treated with 100µM CDCA or 2 µM OCA. (H) Relative SPTLC mRNA level in mouse and human primary cultured hepatocytes after supplementation with 2µM OCA. Bars represent mean ± SD; N=3. Statistical significance was assessed by one-way ANOVA followed by Tukey’s correction (\**P* < 0.05, \*\**P* < 0.01, \*\*\**P* < 0.001, \*\*\*\**P* < 0.0001).

Next, we sought to define if bile acid/FXR signaling pathway may regulate *de novo* SL synthesis in hepatic tissues. Functionally related genes often exhibit coordinated spatiotemporal expression [65]. We thus assessed correlation between SPTLC isoforms and FXR expression in non-tumoral liver tissues from patients with benign liver lesions from a previous cohort [66]. qRT-PCR revealed strong transcriptional correlations between FXR and SPTLC2 or SPTLC3 (Fig. 2C). Similarly, *APOA1*, a gene regulated by the bile acid/FXR signaling pathway [49, 67] also showed a positive correlation with FXR (Fig. 2C). Given the observed transcriptional correlations, we investigated whether FXR activation influences SPTLC gene expression, using Huh7 cells [68–70].

In humans, chenodeoxycholic acid (CDCA) (Fig. 2D) is the most potent endogenous FXR ligand (EC_50_ = 10 μM). Interestingly, CDCA supplementation to Huh7 cells did not affect SPTLC1 or SPTLC2 mRNA levels but significantly reduced SPTLC3 mRNA levels in a dose-dependent manner (Fig. 2E). Obeticholic acid (OCA) is an ethyl derivative of CDCA (Fig. 2D) and a relatively strong FXR agonist (EC_50_ = 0.1 μM) [71]. OCA produced a similar degree of repression of *SPTLC3* transcription at 1/100^th^ of the CDCA extracellular concentration (Fig 2F), supporting bile acid/FXR mediated regulatory effect. Notably, under the same conditions, neither CDCA nor OCA supplementation altered ORMDL expression in Huh7 cells (Fig. 2G). The differential regulation of SPTLC genes by bile acids was further confirmed in human and mouse-derived primary hepatocytes. In both cells lines, OCA supplementation led to a significant and specific reduction in SPTLC3 expression (Fig. 2H). Together these data suggest that among SPT subunits, bile acids specifically regulate *SPTLC3* in hepatocytes.

### FXR binds cis-acting sequence elements in *SPTLC3* promoter

FXR serves as the primary bile acid receptor in mammals [54], mediating both direct and indirect transcriptional repression of its target genes [49, 52]. Chromatin immunoprecipitation (ChIP) assays were conducted to confirm (or rule out) direct FXR binding at the *SPTLC3* locus. In *silico* analysis of the contiguous 2000 bp human *SPTLC3* promoter identified two possible FXR binding half sites, ^1984−^ACTTCAA^−1979^ (FXR *cis*-element 1; reverse complement) and ^593−^AGGTCA^−588^ (FXR *cis*-element 2) upstream of the transcriptional start site (TSS) (Fig. 3A) and (Supplementary sequence file 2). The RT-qPCR of ChIP eluates using site-specific primers showed enrichment of *cis*-element 2 but not *cis*-element 1 in the immunoprecipitated DNA (Fig. 3B). Reduction in FXR expression with siRNA abrogated the immuno-precipitation of *cis*-element 2 (Fig. 3B and C). Additionally, OCA supplementation in Huh7 cells increased FXR binding at the *cis*-element 2, as evidenced by significant enrichment of this target sequence in the bile acid treated cells (Fig. 3D). Therefore, *cis*-element 2 appeared to be the site for FXR binding in the SPTLC3 promoter. We investigated whether the enhanced enrichment of the FXR binding sequence after OCA supplementation was due to the presumed cytoplasm-to-nuclear translocation of the activated FXR receptor. To address this question comprehensively, cytoplasmic and nuclear fractions were separated by differential centrifugation. Western blot analysis of these fractions showed that, irrespective of treatment or OCA dose, FXR was largely present in the nuclear fractions (Fig. 3E). A low FXR presence was observed in the cytoplasmic fractions, that were absent in siRNA-transfected cells (Fig. 3E). However, these cytoplasmic FXR protein bands did not change between vehicle and OCA supplemented cells. These data may indicate that intra-nuclear activation of FXR may account for its enrichment at the SPTLC3 promoter element 2 in OCA treated cells.

**Figure 3:**
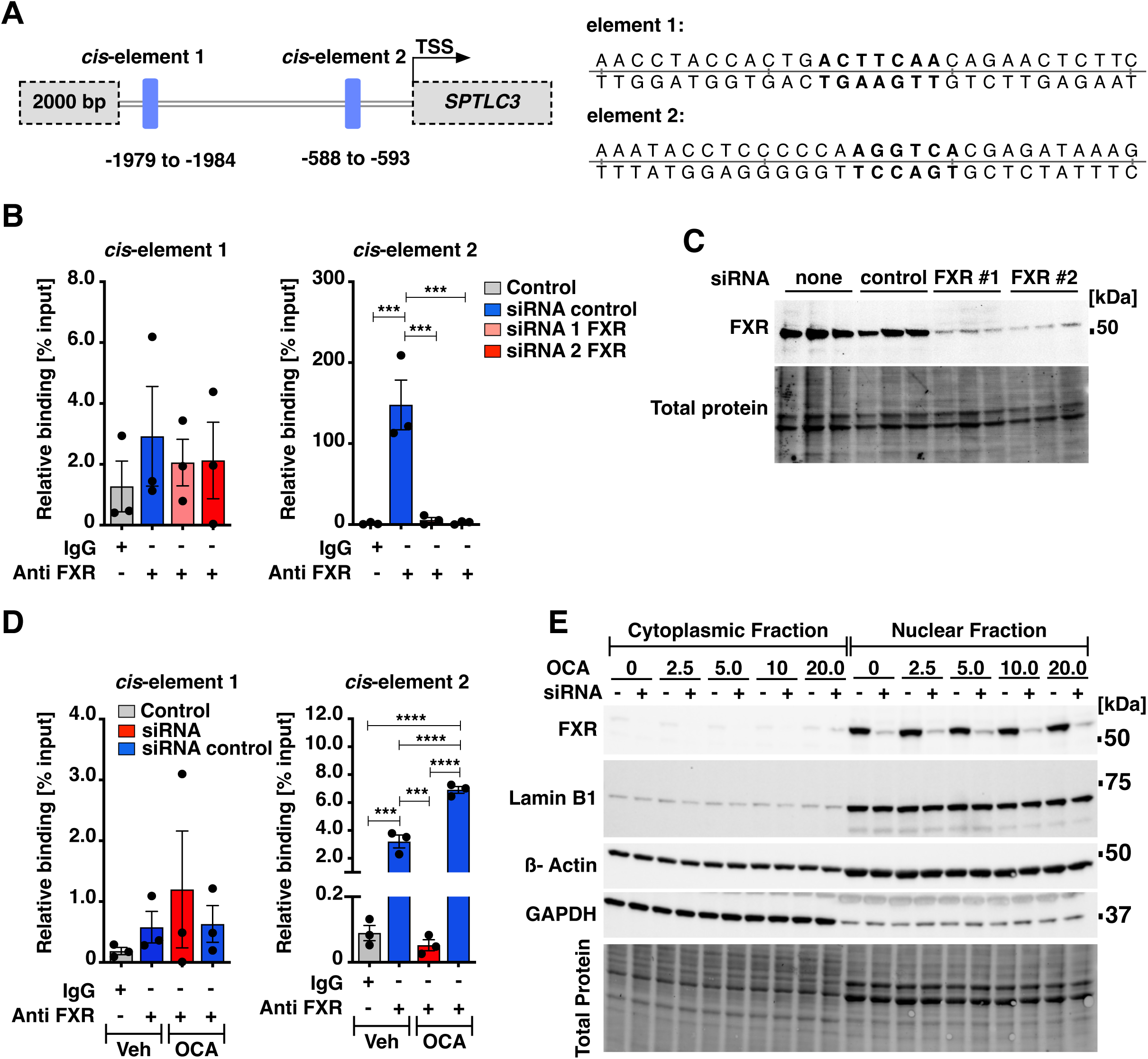
FXR directly binds SPTLC3 promoter. (A) In *silico* identified putative FXR binding sites, *cis*-element 1 and 2, in the proximal SPTLC3 promoter. (B) Chromatin-immunoprecipitation (ChIP) in Huh7 cells using FXR antibody. (C) Western blot of FXR after silencing with two different siRNAs. (D) ChIP assay in Huh7 cells after OCA supplementation. (E) FXR cytoplasmic and nuclear localization in Huh7 cells under the indicated experimental conditions. Bars represent mean ± SD; N=3 (B and D). Statistical significance was assessed by one-way ANOVA followed by Tukey’s correction (\**P* < 0.05, \*\**P* < 0.01, \*\*\**P* < 0.001, \*\*\*\**P* < 0.0001).

To test FXR specificity in bile acid-mediated repression of SPTLC3, Huh7 cells were subjected to FXR KD by siRNA followed by expression analysis. FXR KD did not affect basal expression of SPTLC genes, but completely abolished OCA mediated *SPTLC3* repression (Fig. 4A). FXR often exerts gene regulation in conjunction with RXRα. Silencing RXRα diminished but didn’t abolish OCA mediated SPTLC3 repression in Huh7 cells (Fig. 4A). We conclude that FXR regulation of SPTLC3 is likely independent of RXRα. Interestingly, the RXRα agonist, 9-*cis* retinoic acid (9-*cis* RA) [72] also induced a dose dependent repression of SPTLC3 expression (Fig. 4B).

**Figure 4:**
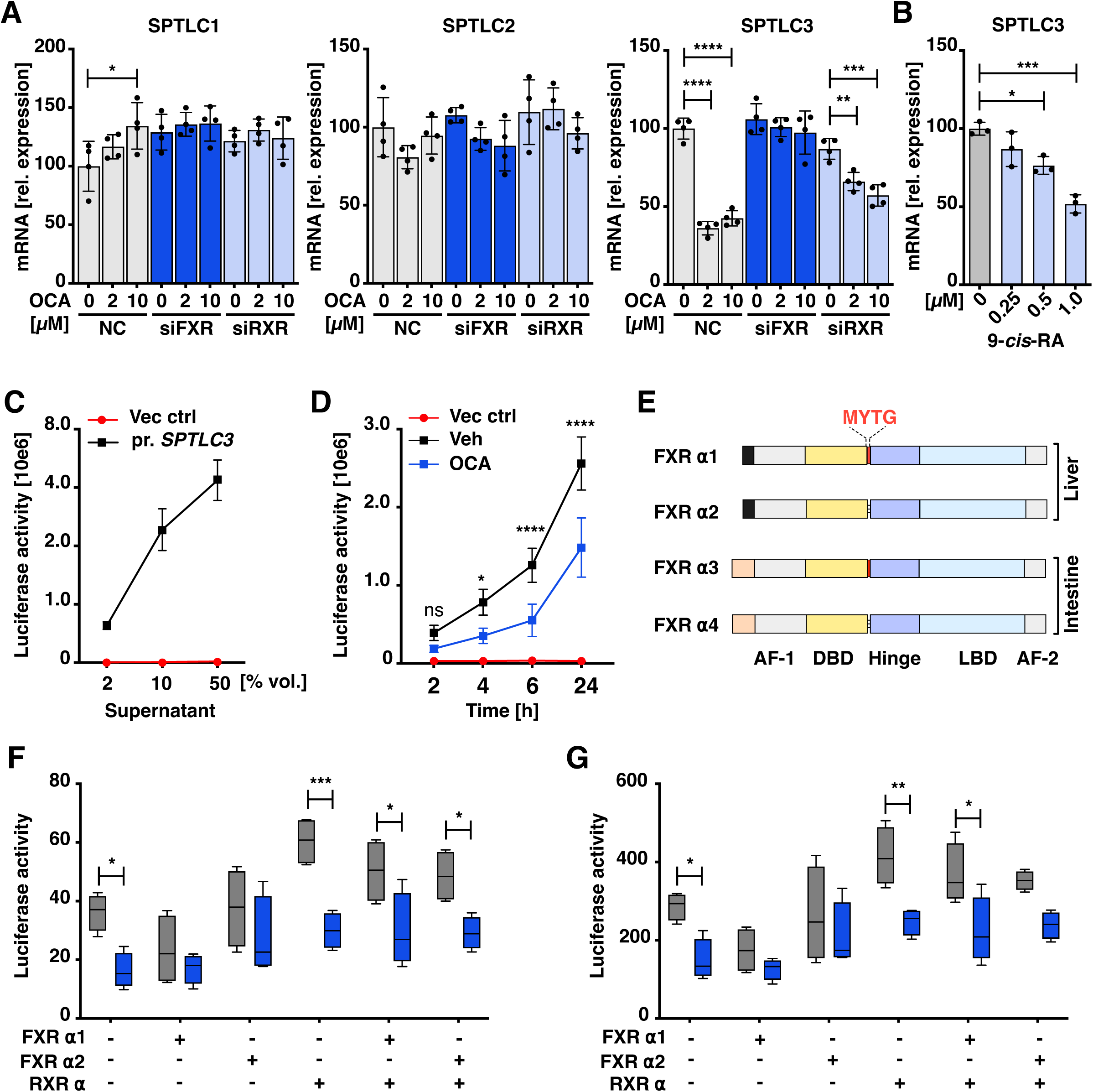
SPTLC3 promoter cis-element 2 is a negative FXR sequence element. (A) Expression of SPTLC1 isoforms in Huh7 cells after silencing of FXR and retinoid x receptor alpha (RXRα). (B) Relative SPTLC3 mRNA levels in Huh7 cells treated with the RXRα agonist, 9 cis-retinoic acid. (C) SPTLC3 promoter and luciferase activity from the culture media of promoter-SPTLC3-*Gaussia* luciferase (Pr.SPTLC3-GLuc) plasmid transfected cells over time. (D) Luciferase activity from media from Pr.SPTLC3-GLuc-transfected and OCA supplemented Huh7 cells. (E) Cartoon depicting domain organization of known human FXR isoforms expressed in liver and intestinal cells. (F-G) Luciferase activity in the media and protein lysates from Pr.SPTLC3-GLuc co-transfected with hepatic FXR isoforms and RXRα. Bars represent mean ± SD; N=3 (B and D). Statistical significance was assessed by one-way ANOVA followed by Tukey’s correction (\**P* < 0.05, \*\**P* < 0.01, \*\*\**P* < 0.001, \*\*\*\**P* < 0.0001).

Furthermore, luciferase assays were used to confirm whether the FXR-responsive cis-element 2 within the SPTLC3 promoter is sufficient to mediate gene repression. To this end, the 1605 bp promoter region flanking the 5’-*SPTLC3* was cloned (Supplementary sequence file 2) upstream of *Gaussia* luciferase (GLuc) coding sequence [73]. First, Huh7 cells were transfected with control and promoter *SPTLC3*-Gluc (pr.*SPTLC3*-GLuc) plasmids. Activity of the secreted GLuc in the growth medium presents as a sensitive readout of the target promoter activity [74]. Only medium fractions from pr.*SPTLC3*-GLuc transfected Huh7 cells showed significant luciferase activity, confirming promoter activity in these cells (Fig. 4C). The GLuc activity also increased over-time in media replenished pr.*SPTLC3*-Gluc-transfected cells only, while OCA supplementation significantly decreased the GLuc activity (Fig. 4D).

Human FXR, due to alternate promoter usage and/or splicing, is expressed as four isoforms, FXRα1-4. These isoforms differ in their N-terminal activation function domain 1 (AF-1) and presence of Met-Tyr-Thr-Gly (MYTG) amino acid extension of the DNA binding domain [44] (Fig. 4E). FXRα1 and FXRα2 are predominantly expressed in liver, while FXRα3 and FXRα4 are primarily expressed in the kidneys and intestines (Fig. 4C) [75]. Since, FXR mediates both isoform specific and non-specific transcriptional regulation [54], we investigated isoform specificity in SPTLC3 regulation. Luciferase assays were performed from media fractions of FXRα1 or FXRα2 and RXRα plasmids transfected Huh7 cells (Fig. 4F). To rule out cellular retention of GLuc protein in OCA supplemented cells, luciferase assays were also performed from cell lysates. Basal GLuc activity was observed to be lower in the media and protein fractions from FXRα1 than the FXRα2 transfected cells (Fig. 4F and G). Interestingly, RXRα co-transfection elevated basal GLuc activity (Fig. 4F and G). The OCA decreased GLuc activity in the media and cells from all plasmid combinations tested (Fig. 4F). However, the strongest relative repression of GLuc activity could be observed in RXRα and RXR/FXR transfected cells. Furthermore, the lack of *SPTLC3* and FXR expression in HEK293 cells or their presence in hepatocytes [6, 76] may suggest FXR driven cell specific SPTLC3 expression. However, FXR, RXRα expression in HEK293 cells did not induce *SPTLC3* expression (Supplementary Fig. 4).

Collectively, these results indicate that FXR represses SPTLC3 transcription in hepatocytes by directly binding to its promoter. The bile acid/FXR mediated regulation is independent of RXRα and likely FXR isoform specific.

### Bile acid-FXR dependent regulation of hepatic *Sptlc3* in mice

Next we assessed FXR-mediated regulation of hepatic *Sptlc3 in vivo* in mice. Orthologous transcription factor binding is often conserved in promoters than in distal regulatory elements [77]. Sequence alignment of the human and mice SPTLC3 proximal promoter regions showed that the FXR binding cis-element 2 but not the non-FXR binding cis-element 1 was conserved between the two species (Supplementary Fig. S5A). However, analysis of publicly available single-cell transcriptomic data from the mouse liver revealed minimal to no *Sptlc3* expression in major embryonic and adult hepatic cell types, including hepatocytes (Supplementary Fig. S5B). In contrast, HFD robustly induces *Sptlc3* expression in mouse liver [32, 34, 35]. To test whether bile acids may counteract *Sptlc3* induction by HFD, female mice were fed either a chow or HFD for 8 weeks, followed by co-supplementation with vehicle or OCA. Consistent with diet-induced metabolic disease, HFD fed mice presented with elevated fasting blood glucose, increased body weight, plasma triacylglycerol (TAG), and total cholesterol, phenotypes that were normalized with OCA (Supplementary Fig. S5C and Supplementary Table 1). Hepatic lipid accumulation in HFD-fed mice was also attenuated by OCA co-supplementation (Fig. 5A). These results confirm the expected metabolic consequences of HFD and bile acid supplementation.

**Figure 5:**
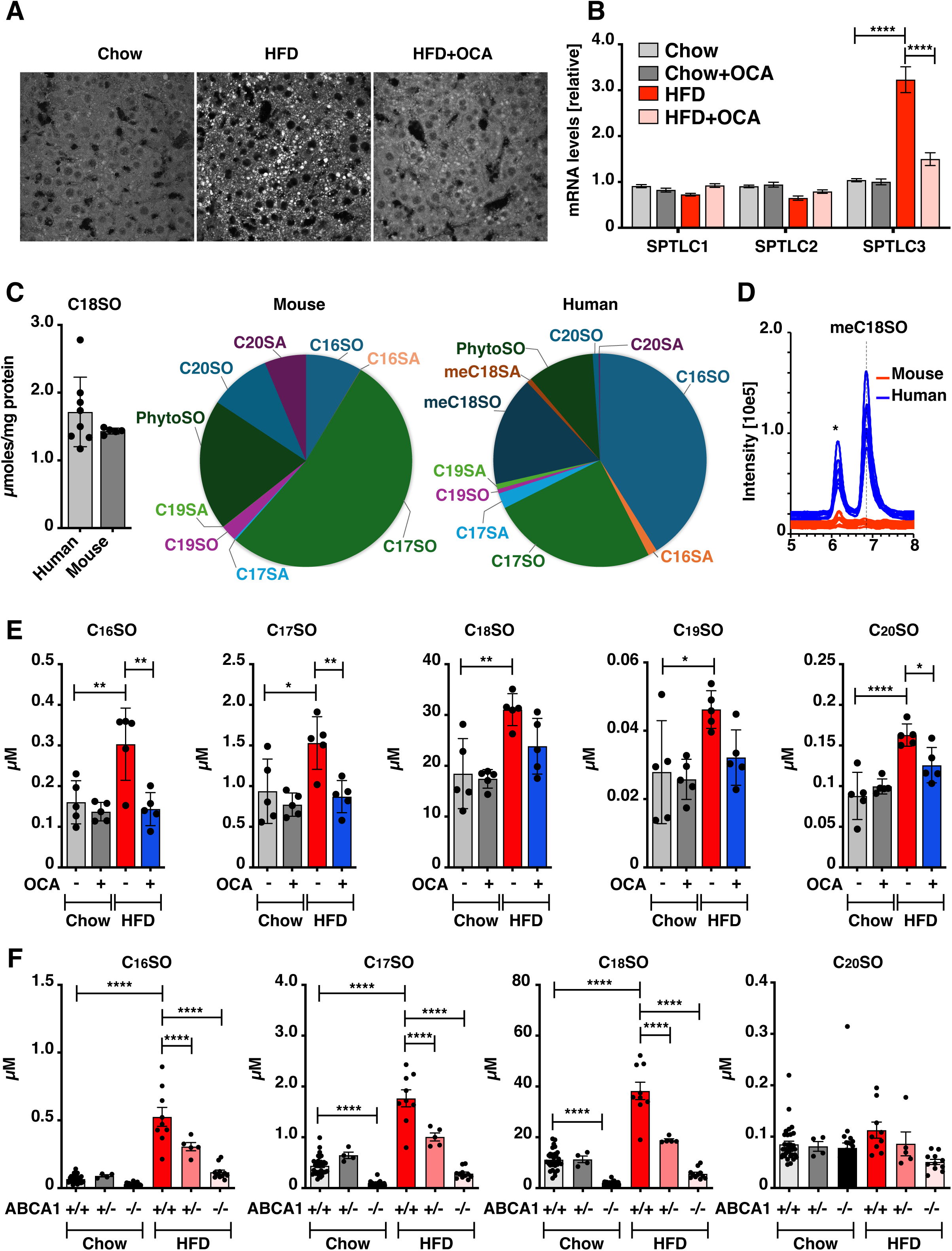
FXR regulates SPTLC3 expression in livers of mice on high fat diet (HFD). (A) Representative BODIPY-stained images illustrating hepatic lipid accumulation (white blobs) in control and HFD fed mice with and without obeticholic acid (OCA) mice. (B) Hepatic SPTLC1, 2 and 3 mRNA levels in mice upon the indicated diet interventions. (C) Levels of sphingosine, C_18_SO (bars) and relative abundance of atypical long chain bases (LCB, pie-charts) in mice and human liver tissue. (D) Comparisons of LC-MS chromatograms showing presence of the omega-3-methylsphingosine (meC_18_SO) in human but not in mouse derived liver lysates. ‘*’ depicts yet unidentified LCB. (E) Plasma LCB profiles from mice on chow and HFD fed and co-supplemented with OCA. (F) Plasma LCB levels from control and hepatocyte-specific ATP binding cassette subfamily protein 1 (Abca1) heterozygous (+/−) and homozygous (−/−) deletion mice, with and without HFD. Bars represent mean ± SD; Statistical significance was assessed by one-way ANOVA followed by Tukey’s correction (\**P* < 0.05, \*\*\**P* < 0.001, \*\*\*\**P* < 0.0001).

Quantification of mRNA levels of SPTLC genes across different groups revealed significant upregulation of hepatic *Sptlc3* expression in response to HFD (Fig. 5B). OCA supplementation with HFD specifically repressed *Sptlc3* to the levels seen in chow fed mice (Fig. 5B).

Next, we examined the metabolic consequences of FXR-mediated repression of *Sptlc3*. First, we conducted a cross-species comparison of hepatic LCB profiles. Tissue homogenates from human livers from our cohort [66] and control mice were analyzed, revealing marked differences in LCB profiles between the species. C_18_SO is the most abundant LCB in both species, with comparable levels in human and mouse liver (Fig. 5C). Among atypical LCBs, C_17_SO is over five times more abundant than C_16_SO in mouse liver (Fig 5C). In contrast, C_16_SO is most prevalent atypical LCB in human liver, being approximately 1.5 fold higher than C_17_SO (Fig 5C). C_20_SO is relatively abundant in mouse than in the human liver. The corresponding saturated LCB followed similar trend in occurrence and abundance (Fig. 5C). Among the isobaric C19 LCBs, straight chain C_19_SO is the least abundant in the livers of either species, while meC_18_SO was detected exclusively in human liver tissues (Fig. 5C and D).

Next, LCB levels were compared among control, HFD, and OCA-supplemented mice groups. Interestingly, hepatic LCB levels remained comparable regardless of the treatment (Supplementary Fig. 5D). However, a similar analysis revealed significantly increased levels of C_16_SO, C_17_SO, C_18_SO, and C_20_SO LCBs in plasma of HFD-fed mice (Fig. 5E). OCA co-supplementation maintained low plasma levels of all LCBs in HFD-fed mice (Fig. 5E). However, the decrease in plasma LCB levels in HFD-OCA-supplemented mice was significant for C_16_SO, C_17_SO, and C_20_SO only. The plasma LCB levels thus correlate with SPTLC3 expression in the mouse liver. Furthermore, the branched meC_18_SO was not detectable in the liver or plasma of mice, even after HFD intervention.

Other than liver, intestines possess significant FXR activity and are a major source of plasma lipids [44]. Therefore, we further sought to confirm the hepatic origin of SPTLC3-derived plasma SL under HFD. ATP binding cassette subfamily protein 1 (ABCA1) facilitates intra- and extra-cellular lipidation of ApoA. ABCA1 loss of function mutations cause Tangier disease (TD), a rare genetic disorder with hypolipoproteinemia in patients and mice models of the disease [78–80]. Plasma samples were analyzed from wild type and mutant mice with hepatocyte specific hetero- and homozygous deletion ABAC1 deletion [80]. Plasma LCBs were reduced in the hetero- and homozygous ABCA1 KO mice compared to homozygous wild type animals on chow diet (Fig. 5F). HFD increased plasma LCB levels in the wild type and mutant mice, but the increase was significantly higher in wild type mice than the mutants (Fig. 5F). Plasma LCB levels decreased progressively from wild-type to heterozygous to homozygous ABAC1 KO mice, indicating a dosage dependent effect of ABCA1 expression. These data suggest hepatic origin of plasma LCBs or SL. These data also suggest that plasma SL may be modulated by lipoprotein assembly factors in the organ.

### MASLD patients accumulate SPTLC3 derived LCB

To evaluate the association between SPTLC3-derived SL and metabolic dysfunction in humans, we quantified plasma LCB profiles in a cohort of MASLD patients (n = 56; 34 males, 22 females) and healthy individuals (n = 33; 17 males, 15 females, 1 unspecified) (Supporting Table 2). As expected, the patients showed increased plasma TAG and cholesterol levels (Supporting Table 2). LCB comparisons between the two groups showed that SPTLC3 derived C_16_SO, C_17_SO, meC_18_SO, and C_20_SO were elevated in MASLD patients (Fig. 6A). C_18_SO levels appeared similar in two groups (Fig. 6A). 1-deoxySL [1-deoxySA (m18:0) and 1-deoxySO (m18:1)], atypical LCB formed when SPT utilizes alanine instead of its usual substrate serine [81], were also elevated in MASLD patients (Fig. 6B). A volcano plot analysis revealed the highest relative increase in meC_18_SO in MASLD patients, followed by C_16_SO (Fig. 6C). Interestingly, phytosphingosine (phytoSO; t18:0) that is the main LCB found in plants and yeast, appeared decreased in the patients (Fig. 6C).

**Figure 6:**
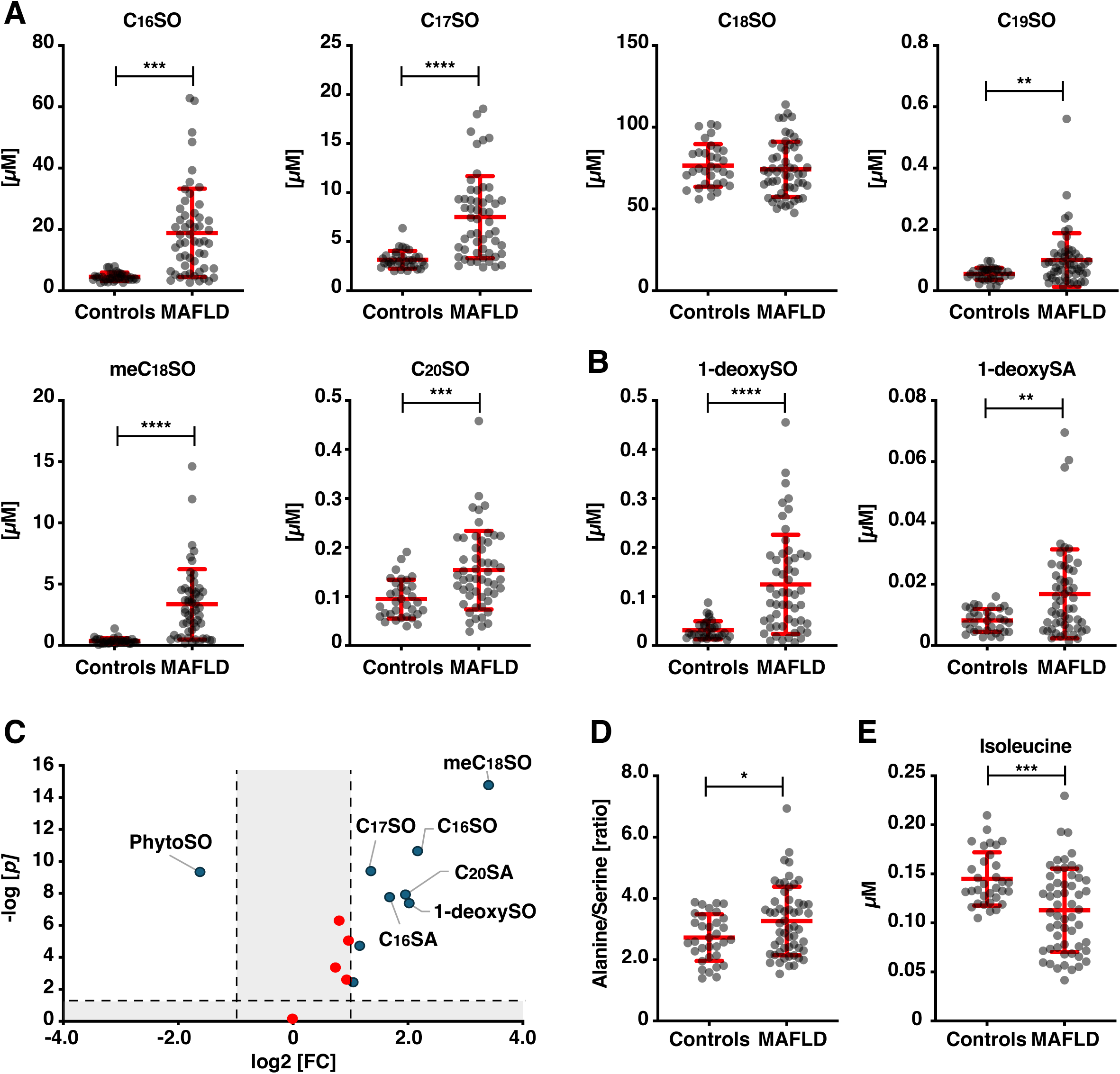
Plasma LCBs in metabolic associated fatty liver disease (MASLD) patients. (A) Plasma levels of sphingosine (C_18_SO) and SPTLC3 derived atypical long chain bases (LCBs) from healthy (N=33) and MASLD patients (N=56). (B) Plasma levels of 1-deoxysphingolipids; 1-deoxysphinganine (1-deoxySA) and 1-deoxysphingosine (1-deoxySO) in the two groups. (C) Volcano plot depicting statistical significance (−log_10_ *P*-value) versus log_2_ fold-change (FC) in LCBs between control and MASLD patients. (D) Ratio of SPT substrate amino acids, alanine and serine in control and MASLD groups. (E) Levels of branched chain amino acid, isoleucine in control and patient plasma. Bar graphs represent mean ± SD (A-B and D-E). Statistical comparisons between groups was performed using two-tailed unpaired Mann-Whitney test (\**P* < 0.05, \*\**P* < 0.01, \*\*\**P* < 0.001, \*\*\*\**P* < 0.0001).

Amino acids such as Ser, Ala and Ilu may modulate LCB formation in human cells. A higher Ala to Ser ratio increases 1-deoxySL formation [18, 82, 83]. Similarly, the branched fatty acid for meC_18_SO synthesis is derived from Ileu, [6]. Therefore, plasma amino acids from control and MASLD patients were quantified by LC-MS. Total Ser and Ala were reduced in the patients (Supplementary Fig. 6A), yet the Ala/Ser ratio increased significantly (Fig. 6A). However, Ile levels were decreased in patients (Fig. 6B), as were Leu and Val (Supplementary Fig. 6B). Therefore, increased SPTLC3 derived plasma LCBs are likely due to increased expression/activity of the enzyme.

It is technically challenging to specify functions of SL derived from individual SPTLC3 generated LCB in isolation. Nevertheless, SPTLC3 deficiency decreases mitochondrial respiration and ROS production in murine cardiomyocytes [40]. Given its human specific presence and significant association with MASLD patient phenotype, we asked whether meC_18_SO may alter mitochondrial function/ROS production, and effect hepatocyte survival in vitro. First, we evaluated the effect of meC_18_SO supplementation on Huh7 cell viability. At low meC_18_SO levels (1 and 5 µM; plasma concentration of the healthy and MASLD patients, respectively), there was only a marginal and non-significant decrease in Huh7 cell viability within 2 h of supplementation. At higher meC_18_SO concentrations (25 µM), cell viability decreased drastically (Fig. 7A). Next, we assessed ROS production in Huh7 cells expressing the redox-sensitive green fluorescent protein, localized to the mitochondria, Mito-roGFP or cytoplasm, Cyto-roGFP [84]. Oxidation-reduction status indicated dose dependent increase of ROS levels in meC_18_SO supplemented Mito-roGFP expressing Huh7 cells (Fig. 7B). Mitochondrial ROS levels already increased at sub-toxic meC_18_SO concentrations (1 and 5 µM) (Fig. 7B). In contrast, fluorescence levels in Cyto-roGFP expressing Huh7 cells increased at high meC_18_SO dose only. We conclude that at concentrations relevant in vivo, meC_18_SO induces ROS production primarily in the mitochondria.

**Figure 7:**
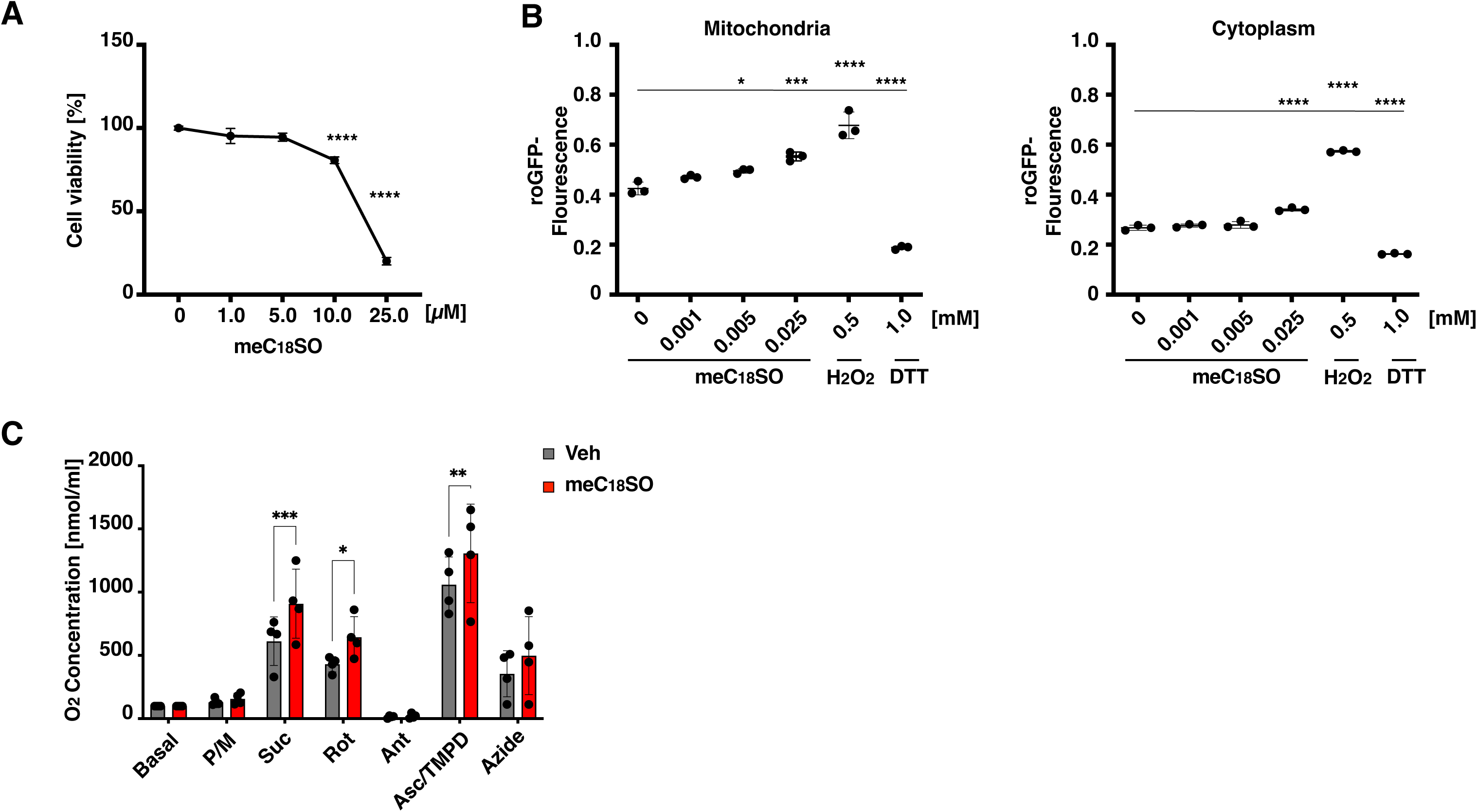
Omega-3-methylsphingosine (meC_18_SO) affects ROS levels and mitochondrial respiration in Huh7 cells. (A) Viability of Huh7 cells supplemented with meC_18_SO. (N=3) (B) Mitochondrial and cytoplasmic oxidative stress/ROS production in Huh7 cells expressing Mito-roGFP and Cyto-roGFP, respectively. (N=3) (C) Mitochondrial respiration in Huh7 cells as measured with O2k-FluorRespirometer (Oroboros) (N=4) Data is represented as mean ± SD. Statistical comparisons were performed using 2-way Anova (A and C), ordinary one-way ANOVA (A) with Bonferroni correction, (\**P* < 0.05, \*\**P* < 0.01, \*\*\**P* < 0.001, \*\*\*\**P* < 0.0001).

Next, we evaluated oxygen consumption rate (OCR) in meC_18_SO supplemented Huh7 cells using O2k-FluorRespirometer with a multiple substrate-uncoupler-inhibitor titration protocol for electron transport chain (ETC) complex’s (CI to IV). CI-substrates, pyruvate and malate, had no effect on the respiration rate (Fig. 7C). However, succinate and ascorbate that are, respectively, the C II and C IV substrates, increased OCR significantly in meC_18_SO compared to the vehicle treated Huh7 cells.

These data indicate that meC_18_SO likely acts as a modulator of mitochondrial function and ROS levels in human hepatocytes.

## Discussion

SPTLC3 expression and activity are linked to dyslipidemia associated hepatic and cardiovascular pathology. Accordingly, identifying factors that regulate SPTLC3 expression and activity remain a key objective with potential pharmacological relevance. In this study, we define two distinct modes of SPTLC3 regulation: post-translational inhibition by ORMDL proteins and transcriptional repression by the bile acid-activated nuclear receptor FXR. These regulatory mechanisms shape hepatic SL output and are associated with metabolic dysfunction in both mouse models and human.

The activity of SPTLC1/SPTLC2 complex is inhibited by ORMDL proteins through interaction with the N-terminal transmembrane domain of SPTLC1 [10, 18]. Consistent with this scaffolding role of SPTLC1, ORMDLs modulate SPTLC3-containing SPT complexes via post-translational feedback inhibition (Fig. 1B-D). Since, C_6_-Cer induced inhibition of SPT independent of the subunit composition. The LCB component of this ceramide analogue is SO. These results suggest that ORMDLs may sense overall SL cellular load in the cells. Additionally, single nucleotide polymorphisms (SNPs) that increase hepatic SPTLC3 expression are associated with elevated hepatic lipid content [27]. Conversely, ORMDL3 expression correlates negatively with body mass index (BMI) or related insulin resistance in humans and mice [24, 85]. These findings raise the possibility that SPTLC3-ORMDL interaction contributes to hepatic lipid homeostasis. Therefore, limiting SL production, for example, by pharmacological induction of ORMDL expression [86] may have therapeutic potential in metabolic conditions.

Mechanism(s) that stimulate SPTLC3 expression in MASLD [32, 35, 87] is not clear. Incoherent cellular responses to lipotoxic stress likely contribute to the upregulation of SPTLC3. However, it is striking that bile acids/FXR regulate SPTLC3 in hepatic cells but not the ubiquitously expressed SPTLC1 or SPTLC2 (Fig 3A-D and Fig. 4A-D). ChIP and luciferase assays confirmed the half FXRE ^593−^AGGTCA^−588^ site at the SPTLC3 promoter as the negative FXR response element. This follows a pattern where FXR independently binds the unusual DNA elements, for example in APOA1, for transcriptional repression of its target gene [49]. However, existence of additional negative FXRE’s within the regulatory or non-coding regions of SPTLC3 cannot be ruled out. Although bile acid profiles differ between humans and mice [88, 89], OCA supplementation repressed SPTLC3 in mice liver (Fig. 5B), supporting FXR-mediated repression in vivo.

SPTLC3 expression in the liver correlated with plasma LCB levels, but not with hepatic LCB content (Fig. 5 and Supplementary table 1). However, plasma LCB levels in both mice and humans mirrored the signature of SPTLC3-expressing cells (Fig. 5C-E; Fig. 6A-C). Also, reduced plasma SL levels in hepatocyte-specific *Abca1* knockout mice strongly support hepatic origin for circulating SPTLC3-derived species (Fig 5F). In our analysis, cholesterol was also increased in plasma but not liver of the HFD-fed mice (Supplementary table 1). These observation likely reflect increased co-efflux of cholesterol and SL from the liver under HFD conditions. Transcriptomic data and qPCR analysis revealed higher SPTLC3 than SPTLC2 expression in human hepatocytes and liver (Supplementary Fig. S3A and Fig. 2C). Since LCBs are obligate precursors of complex SL, hepatic expression of SPTLC3 highlights its role in diversifying circulating SL pool from the outset. Additionally, increase in SPTLC3 specific products over the canonical C_18_SO in humans, may induce changes downstream of the SL pathway. This remodeling may underlie disease-associated lipid profiles. FXR-mediated repression of *SPTLC3* in liver targets the earliest step in sphingolipid biosynthesis, inducing a metabolic shift in circulating SL.

While SPTLC3 derived products discriminated effectively between MASLD and healthy states, meC_18_SO appeared the most significantly elevated in the patient plasma (Fig. 6C). Prior studies have associated C19-LCB-based SLs with SPTLC3 and CVD [30, 38]. Our analysis revealed that meC_18_SO is present at markedly higher levels than the straight-chain C_19_SO in human liver and plasma (Fig. 5C, D and Fig. 6A-C). Therefore, meC_18_SO based SL are likely indicators for metabolic dysfunction. Further verification in independent patient cohorts will be needed to ascertain such facts.

The physiological relevance of increased SPTLC3 derived SL in MASLD remains unclear. SLs containing atypical LCBs are expected to alter membrane properties, signaling, and cellular function, through modulation of membrane organization [34, 90]. C_16_SO-based SL modulate mitochondrial function in ischemic heart tissue at C I of the electron transport chain [34, 40]. The meC_18_SO decreased Huh7 cell viability and elevated ROS production (Fig. 7A and B). However, in contrast to C_16_SO, meC_18_SO enhanced CII and C1V function (Fig. 7). Thus, structural variations in LCB may contribute to the broader and possibly species-specific effects of SPTLC3-derived SLs in humans. Additionally, C1 and CIII primarily contribute to ROS within cells. CII is known to contribute to mitochondrial ROS production under pathological conditions, such as ischemia-reperfusion injury by driving Reverse Electron Transport (RET) at CI [91]. Although more work would be need to ascertain its function, these data provide the first clues to the physiological relevance of this sphingolipid in metabolic disease.

FXR agonists are under clinical investigation for MASLD treatment [43]. In light of recent studies highlighting the pleiotropic roles of SPTLC3 in the liver and heart [34, 39, 40], it is tempting to speculate that bile acid-based therapies may confer benefits, at least partially, through SPTLC3 modulation. As SPTLC3 expression is influenced by dietary stimuli and genetic factors, targeting the enzyme or its upstream regulators may offer therapeutic benefit in metabolic disease.

## Materials and methods

### Cell lines

HEK293 cells were cultured in Dulbecco’s Medium (DMEM, Sigma-Aldrich, St. Louis, MO, USA) with 10% fetal calf serum (FCS). Huh7 cells were maintained in RPMI 1640 medium supplemented with 10% (FCS), 100 U/mL penicillin, 100 μg/mL streptomycin at 37°C in a humidified atmosphere of 5% CO_2_.

### Isotope labelling assay

L-serine labelling assay and SPT activity measurements were performed as described [61]. HEK293 cells were grown to 70% confluence in DMEM growth media. For labelling, the media was exchanged to L-serine free DMEM (Genaxxon Bioscience, Ulm, Germany) supplemented with 10% FBS, 1% P/S and isotope-labelled D_3_-^15^N-L-serine (1 mM) (Cambridge Isotope Laboratories, MA, USA). Cells were grown in the labelling media for 16 hours. For labelling after siRNA knockdown of FXR (below), cells were labelled after 48 hours post transfection for the same time. Myriocin and OCA were added together with the labelling media. For lipid analysis, cells were harvested on ice and frozen after counting (Z2 Coulter counter, Beckman Coulter, CA, USA).

### Glucose and lipid analysis

Blood glucose was assessed via tail tip blood sampling using a glucometer (mylife Unio; Ypsomed Distribution AG, Burgdorf, Switzerland). Cholesterol and triglyceride (TAG) levels from mouse serum were measured using the Amplex Red cholesterol assay kit (A12216; Invitrogen) and the triglyceride assay kit (ETGA-200, EnzyChrom), respectively. Long-chain base analysis was performed as described [59]. Briefly, lipids from frozen cell pellets or plasma (50 µL) were extracted with methanol. Hydrolyzed lipids were resuspended in 70 µL reconstitution buffer (70% methanol, 10 mM ammonium acetate, pH 8.5). 10 µl of the resuspended LCBs were injected into QTRAP 6500+ LC-MS/MS System (Sciex) and lipids separated via a reverse phase C18 column (Uptispere 120 Å, 5 μm, 125×2 mm, Interchim, France). Chromatography and electrospray ionization method were essentially as described [6, 59].

### Hepatic sterol analysis

Three-hundred µg of liver homogenate was mixed with 500 µL of methanol:methyl-tert-buthyl-ether:dichloromethane (4:3:3, MMD), shaken for 1h, at 37°C, and then centrifuged at 16,000g for 5 min at room temperature. The supernatant was dried under a nitrogen stream at room temperature. The dried lipids were hydrolyzed at 65°C for 1 h with 500 µL of 1 M KOH in methanol, mixed with 700 µL of n-heptane, and then centrifuged at 16,000g for 5 min at room temperature. The upper phase was collected, while the lower phase was mixed with 700 µL of n-heptane and centrifuged at 16,000g for 5 min at room temperature, to extract the remaining lipids. The upper phases were combined, dried under a nitrogen stream at room temperature and solubilized in chloroform. Samples were loaded on a HPTLC Silica gel 60 plate with a concentrating zone (Merck KGaA, Darmstadt, Germany) using an automated Camag TLC sampler ATS4 (Muttenz, Switzerland) and separated by one-dimensional thin layer chromatography (TLC). Cholesterol was resolved in n-hexane:n-heptane:diethyl ether:acetic acid (62.4:18.3:18.3:1). Staining was performed in 9.6% orthophosphoric acid (v/v) and 3% copper acetate (w/v), and then the lipids were charred at 130°C for 30 min. Bands were scanned at 366 nm in a Camag TLC Scanner 3 (Muttenz, Switzerland), and absolute quantification was performed from a serial dilution of cholesterol resolved in parallel. The values were then normalized for the protein content.

### Amino acid analysis

Amino acids were quantified from 10 µL of plasma with 180 µL ice-cold methanol containing 1 nmol of stable isotope labelled amino acids (Cambridge Isotope Laboratories, MSK-A2-1.2) as described earlier [18]. Samples were incubated at −20°C for 30 min followed by centrifugation at 4°C, 14,000 x g for 10 minutes. The supernatant was transferred to a fresh tube and dried under a nitrogen stream. Dried samples were stored at −20°C until analysis. For analysis, samples were re-constituted in 100 µL of 0.1 % acetic acid. Amino acids were separated via a reverse-phase C18 column (EC 250/2 NUCLEOSIL 100-3 C18HD, L=250 mm, ID: 2 mm; Macherey-Nagel). Sample (5 µL) were subjected to liquid chromatography coupled multiple reaction monitoring mass spectrometry using a QTRAP 6500+ LC-MS/MS-MS System (Sciex). Buffer systems used were (A) 0.1% formic acid in water and (B) acetonitrile (100%). The amino acids were chromatographed isocratically with solvent A for 5 minutes, followed by a linear gradient to 50% solvent B over two minutes. Then, the column was washed with 80% solvent B prior equilibration with 100% solvent A. The flow rate was held constant at 0.2 mL/min. Sample ionization was achieved via electrospray ionization in positive ion mode. Quantification was performed using MultiQuant (2.1) software (SCIEX).

### Transient transfection with siRNA

Huh7 cells were transfected in 12-well plates at 80% confluency. For each transfection, 5 μL of TransIT-TKO transfection reagent was mixed with 120 μL of serum-free OptiMEM, followed by the addition of either the SMARTpool siRNA targeting FXR or scrambled siRNA at a final concentration of 25 nM. After incubation at room temperature for 10 minutes, the siRNA mixtures were added to 600 μL RPMI 1640 medium supplemented with 10% FBS in each well. Eighteen hours after transfection, the cells were subjected to OCA. Cellular RNAs were extracted using Trizol reagent and subjected to real-time PCR, as described below.

### Animals

Female C57BL/6J mice were randomly assigned to an HFD (D12331; Provimi Kliba, Kaiseraugst, Switzerland) or a chow diet (D12329; Provimi Kliba) for 16 weeks. In a separate experiment, after 8 weeks of high-fat diet, half of the obese mice were given obeticholic acid (OCA) mixed in the food (25 mg/kg, MedKoo Biosciences, NC 27560). Finally, mice were divided into four groups of five animals each: chow, HFD, chow-OCA, and HFD-OCA. Liver from each animal was used for RNA extraction and cDNA generation for RT-PCR analysis. Plasma from C57BL/6J control and hepatocyte-specific Abca1 heterozygous and homozygous knockout mice [80], fed a chow or high-fat diet for 8 weeks [92], was used for LCB measurements.

### Isolation of RNA from liver tissue and cells and quantification of transcript levels

Total RNA was prepared using standard Trizol extraction (Invitrogen, Waltham, MA). Two µg of total RNA were reverse transcribed using random primers and Superscript II enzyme (Invitrogen, Carlsbad, CA). First-strand complementary DNA was used as the template for real-time polymerase chain reaction analysis with TaqMan master mix (Applied Biosystems, Foster City, CA). For the real-time PCR of mouse liver SPTs, the Light Cycler 480 SYBR Green Master kit (Roche, Switzerland) was used. The following primer were: SPTLC1 forward (5’-AGGGTTCTATGGCACATTTGATG-3’), SPTLC1 reverse (5’-TGGCTTCTTCGGTCTTCATAAAC-3’); SPTLC2 forward (5’-CAAAGAGCTTCGGTGCTTCAG-3’), SPTLC2 reverse (5’-GAATGTGTGCGCAGGTAGTCTATC-3’); SPTLC3 forward (5’-TTTGGACTGGACCCTGAAGATATTG-3’), SPTLC3 reverse (5’-TGACTGCATCCGTAAATAATCCACA-3’). Gene expression values were calculated based on the ΔΔCt method. Data were calculated and expressed relative to levels of RNA for the housekeeping gene hypoxanthine phosphoribosyltransferase or β-actin.

### BODIPY staining of mouse liver tissue and microscopy

BODIPY staining was done as described previously [93]. Briefly, the lipid droplet staining was done by BODIPY 493/503 (D-3922; Invitrogen) on liver cryosections. Images were taken by SP8 confocal microscope and analyzed.

### Isolation of primary cultured mouse hepatocytes

Primary cultures of hepatocytes were isolated from female C57BL/6J mice. After a midline incision, a sterile cannula was inserted through the right ventricle and pre-perfusion was performed at 37°C with pre-perfusion buffer (0.5 mM EGTA, 20 mM Hepes in Hanks’ balanced salt solution, pH 7.4) for 10 minutes. The pre-perfusion buffer was then replaced with perfusion buffer (20 mM NaHCO3, 0.5 mg/mL BSA, 6.7 mM CaCl2, 100 U/mL type 2 collagenase, in Hanks’ balanced salt solution, pH 7.4) for 7 minutes. The perfused liver was excised, rinsed in ice-cold William’s medium E with 10% FCS, 2 mM L-glutamine, 2.5 mU/mL insulin, 1 μM dexamethasone and gently disaggregated. After centrifugation, cells were counted, tested for viability, and cultured at 37°C. After 3 hours of incubation, WME-a medium was replaced with WME-b medium (Williams medium E with 10% FCS, 2 mM l-glutamine, 0.25 mU/mL insulin, 0.1 μM dexamethasone).

### Isolation of primary cultured human hepatocytes

Primary human hepatocytes were prepared as previously described [94]. Cells seeded in six-well plates in hepatocyte maintenance medium supplemented with UltraGlutamine for approximately 5 hours before further treatment procedures. Primary human hepatocytes were cultured at 37°C in a humidified atmosphere containing 5% CO_2_.

### Chromatin-Immunoprecipitation (ChIP)

Huh7 cells were transfected with control or NR1H4 siRNA (Qiagen, #1027418), fixed in 1% formaldehyde for 12 min, and quenched with 0.125 M glycine for 5 min at room temperature. After two PBS washes, cells were lysed and nuclei isolated by Dounce homogenization (12 strokes), followed by vortexing on ice and centrifugation (2,500 × g, 10 min, 4 °C). Nuclear pellets were lysed in SDS lysis buffer (50 mM Tris-HCl pH 8.0, 10 mM EDTA, 1% SDS, protease inhibitors) and sonicated (10 cycles, 30 s on/off, 4 °C). Chromatin was cleared (18,000 × g, 20 min), and input DNA (10%) was reserved. Remaining lysates were diluted 1:10 in ChIP dilution buffer and precleared with magnetic beads (Thermo Fisher, Pierce) for 45 min at 4 °C. Samples were incubated overnight at 4 °C with 2 µg anti-FXRα or control IgG. Immune complexes were captured with magnetic beads for 2 h at room temperature and washed twice with low salt, high salt, LiCl, and TE buffers. Chromatin was eluted in ChIP elution buffer (0.1 M NaHCO₃, 1% SDS), reverse crosslinked overnight at 65 °C in 200 mM NaCl, and treated with RNase A and proteinase K. DNA was purified (Meridian Bioscience, #BIO-52066) and analyzed by qPCR using primers, Site 1 forward primer 5’-TTCCTTGTGAGAACCAAGGACT-3’, Site 1 reverse primer, 5’-CATCAGGCACTGCTGGTATGT-3’, site 2 forward primer, 5’-GACCCTGCAGTCATCAACCA-3’, site 2 reverse primer, 5’-GACCCTGCAGTCATCAACCA-3’.

### Luciferase reporter assay

Transcriptional regulation analyses were evaluated using a Promoter SPTLC3-Gaussia-Luciferase reporter plasmid construct (GeneCopoeia, Rockville, MD, USA). Huh7 cells (25, 000 cells per well) were cultured in 24-well plates, transfected with vector control or the reporter plasmids (200 ng per well) alone or with FXR (25 ng and RXRα (25 ng) plasmids. Cells were supplemented with vehicle or OCA, 24 hours post-transfection. Media supernatant or protein lysates were then analyzed for luciferase activity using the luciferase reporter assay system (Promega, E1960), according to the manufacturer’s protocol.

### Oxygen consumption rate (OCR) measurement

Huh7 cell cultures grown overnight in 10 cm plates were treated with vehicle or meC_18_SO for 2 h. Cells were harvested by trypsinization followed by centrifugation (180 rcf, 5 min, RT). Cells (1.0e6) were then resuspended in 2 ml of mitochondrial respiration buffer (MiR05). Oxygen consumption was assessed in the O2k-FluorRespirometer (Oroboros Instruments GmbH, Innsbruck, Austria) using a multiple substrate-uncoupler-inhibitor titration (SUIT) protocol. A pre-calibrated 2-mL chamber was filled with mitochondrial respiration buffer MiR05 and pre-warmed at 37°C. A mini-stirrer was inserted in the chamber and the stirring speed set at 750 rpm. Cells were permeabilized with digitonin (1.25 µg/ml) and then supplemented with Complex I substrates, pyruvate/malate (5 mmol/L, and 2 mmol/L, respectively). Oxygen consumption was activated by ADP (2.5 mmol/L). This was followed by sequential additions of complex II substrate succinate (10 mmol/L) and Complex I inhibitor rotenone (0.5 μmol/L). Subsequently, complex IV substrate, *N*,*N*,*N*′,*N*′-tetramethyl-*p*-phenylenediamine (TMPD) coupled with the TMPD-reduction regeneration system ascorbate (0.5 mmol/L and 2 mmol/L, respectively) were added, followed by complex IV inhibitor sodium azide (70 mmol/L). This allowed determination of sensitive rates of Oxphos using complex I, II, and IV substrates, respectively. OCR was acquired in real-time using the DatLab 7 software. OCR values were corrected for the O_2_ background value (noise) and expressed as pmol of O_2_·sec^−1^·ml^−1^. For each titration, the average value from a one-minute interval at stable OCR was used for data analysis.

### Assessment of oxidative stress in vitro

Plasmids encoding reduction-oxidation sensitive green fluorescent protein (roGFP) localized to the cytoplasm (Cyto-roGFP) or mitochondrial matrix (Mito-roGFP) (AddGene plasmids #49435 and #49437, respectively) were used [84]. Chimeric inserts were sub-cloned into the pcDNA^TM^3.1 ^(+)^ expression vector (Invitrogen, Carlsbad, California, USA) by restriction-ligation cloning using KpnI and NotI restriction sites. Plasmid DNA (1 µg) was transfected into Huh7 cells in six-well plates and harvested by trypsinization 24 h post-transfection, resuspended in PBS and treated for two hours with vehicle or omega-3-methylsphingosine. H_2_O_2_ and DTT (500 µmol/L and 1 mmol/L, respectively) treatments were used as positive controls for oxidized and reduced states of the sensors. Fluorescent signals were then analyzed in a plate reader by sequential excitation with 405 and 475 nm followed by measurement of fluorescence emission at 510 nm. All measurements were conducted in PBS. Data are presented as the ratio of fluorescence after excitation with 405 nm to 475 nm, where a high ratio denotes sensor oxidation and a low ratio denotes sensor reduction [84].

### Study approvals

Animal intervention conformed to Swiss animal protection laws and were approved by the Cantonal Veterinary Office (ZH145/17). Hepatocyte isolation from mice was approved by the Cantonal Veterinary Office (ZH247/15). Human liver tissue for cell isolation was obtained from the charitable state-controlled Human Tissue and Cell Research (HTCR) Foundation, with informed patient consent and approved by the local Ethics Committee. The human study was conducted in accordance with the Declaration of Helsinki guidelines regarding ethical principles for medical research involving human subjects. All patients provided written informed consent, and the study protocol was approved by the Scientific Ethical Committee of Shandong University of TCM, Jinan, China, where the patients were based (license number SDUTCM2010034). The human samples used for figure 2C and 5C were taken from a cohort described in reference [66] (W17_133#17.153). The mouse samples used for figure 5F belongs to experiments described in reference [92].

### Statistical analysis

Data were compared by one-way ANOVA followed by the Tukey’s test. Values of P<0.05 were considered statistically significant. Statistical analyses were performed with GraphPad Prism 8.0 (GraphPad Software, Inc., San Diego, CA).

## Supporting information

Suppementary Table 2

Supplementary sequence 1

Supplementary sequence 2

## Data Availability Statement

All data discussed in this study are included in the main text and supplementary Appendix.

## Acknowledgements

We acknowledge the Human Tissue and Cell Research (HTCR) Foundation for making human tissue available for research. The plasmid constructs for FXR isoforms [50] were a kindly provided by Saskia W. C. van Mil (UMC Utrecht).

## Funding

This work was support by grants from *Schweiz Stiftung für die Erforschung der Muskelkrankheiten* (FSRMM) and the EMPIRIS foundation to Museer A. Lone. Swiss National Science Foundation (SNSF) grants to Gerd A. Kullak-Ublick (#310030_175639), Arnold von Eckardstein (31003A-160126, 310030-185109), Thorsten Hornemann (310030_215134), European Joint Program on Rare Diseases (SNF 32ER30_187505) to Thorsten Hornemann, University of Zurich (Forschungskredit, FK-20-037) to Grigorious Panteloglou. Imme Garrelfs received financial support from the *Amsterdam Universiteits Fonds*, the *Fundatie van Renswoude*, and the *Bekker-la Bastide-Fonds*. Francesca Barone is supported by Next Generation EU, in the context of the National Recovery and Resilience Plan, Investment PE8 – Project Age-It: “Ageing Well in an Ageing Society”. This resource was co-financed by the Next Generation EU [DM 1557 11.10.2022]. The views and opinions expressed are only those of the authors and do not necessarily reflect those of the European Union or the European Commission. Neither the European Union nor the European Commission can be held responsible for them.

## Figure legends (supplementary)

**Figure S1: Sphingolipid (SL) biosynthesis pathway in mammalian cells**

(A) Canonical SL biosynthesis starts with condensation of palmitoyl-CoA and L-serine by serine-palmitoyltransferase (SPT) to form long chain base (LCB). *N*-acylation of the LCB by dihydroceramide synthases (CERS1-6) results in the formation of dihydroceramides. Subsequent desaturation of the LCB backbone by the dihydroceramide desaturase, DEGS1 forms ceramides. Ceramides are converted to complex SLs, sphingomyelin and glycosphingolipids through modification on the 1-hydroxyl group. Turnover of canonical SL occurs through Sphingosine-1-phosphate.

(B) Substrate specificity of SPTLC2 and SPTLC3 containing SPT enzyme and their modulation by ancillary ssSPTa/b units. Cartoon is generated using BioRender.com.

**Figure S2: Long chain base forms in SPTLC3 expressing HEK293 cells**

(A) **The** *de novo* synthesized LCBs in designated HEK293 cells. SPTLC3 induces formation of atypical short (C_16_SO), Odd (C_17_SO), omega-3-methylsphingosine (meC_18_SO), and C_20_SO.

(B) C_6_-Ceramide induced dose dependent inhibition of *de novo* LCB synthesis in SPTLC3 expressing and control HEK293 cells.

**Figure S3: Expression of SPT enzymatic and regulatory subunits in human liver cells.**

(A) Single cell transcriptomic data showing SPTLC1-3 and ORMDL1-3 expression in human liver cells.

(B) Relative expression of these genes from RNA-Seq data in Huh7 cells.

**Figure S4: Ectopic expression of FXR isoforms and RXRα in HEK293 cells.**

Relative expression and mRNA levels of FXR isoforms, RXR, and SPTLC3, in HEK293 cells transfected with FXR isoforms and/or RXR.

**Figure S5: SPTLC3 expression in mice liver cells its metabolic effects.**

(A) Sequence alignment and conservation of FXR site between mouse and human proximal SPTLC3 promoter.

(B) Single cell transcriptomic data showing expression of SPTLC1-3 and ORMDL1-3 genes in liver cells of mice.

(C) Fasting glucose levels in chow and HFD diet-fed mice supplemented with OCA.

(D) LCB profiles in livers of mice fed a chow or HFD, with and without obeticholic acid (OCA)

**Figure S6: SPT linked amino acid levels in metabolic associated fatty live disease (MASLD) patients**

(A) Plasma levels of alanine and serine and (B) branched chain amino acids in control and MASLD patients.

## Notes

### Competing Interest Statement

The authors have declared no competing interest.

