## Supplementary Fig for "Dyslipidemic SPTLC3 Integrates Bile Acid-FXR Signaling with Sphingolipid Remodeling in MASLD"

**Supplementary data:**

**Figures S1 to S6**

**Supplementary Table 1 and 2**

**Supplementary sequence files 1 and 2**

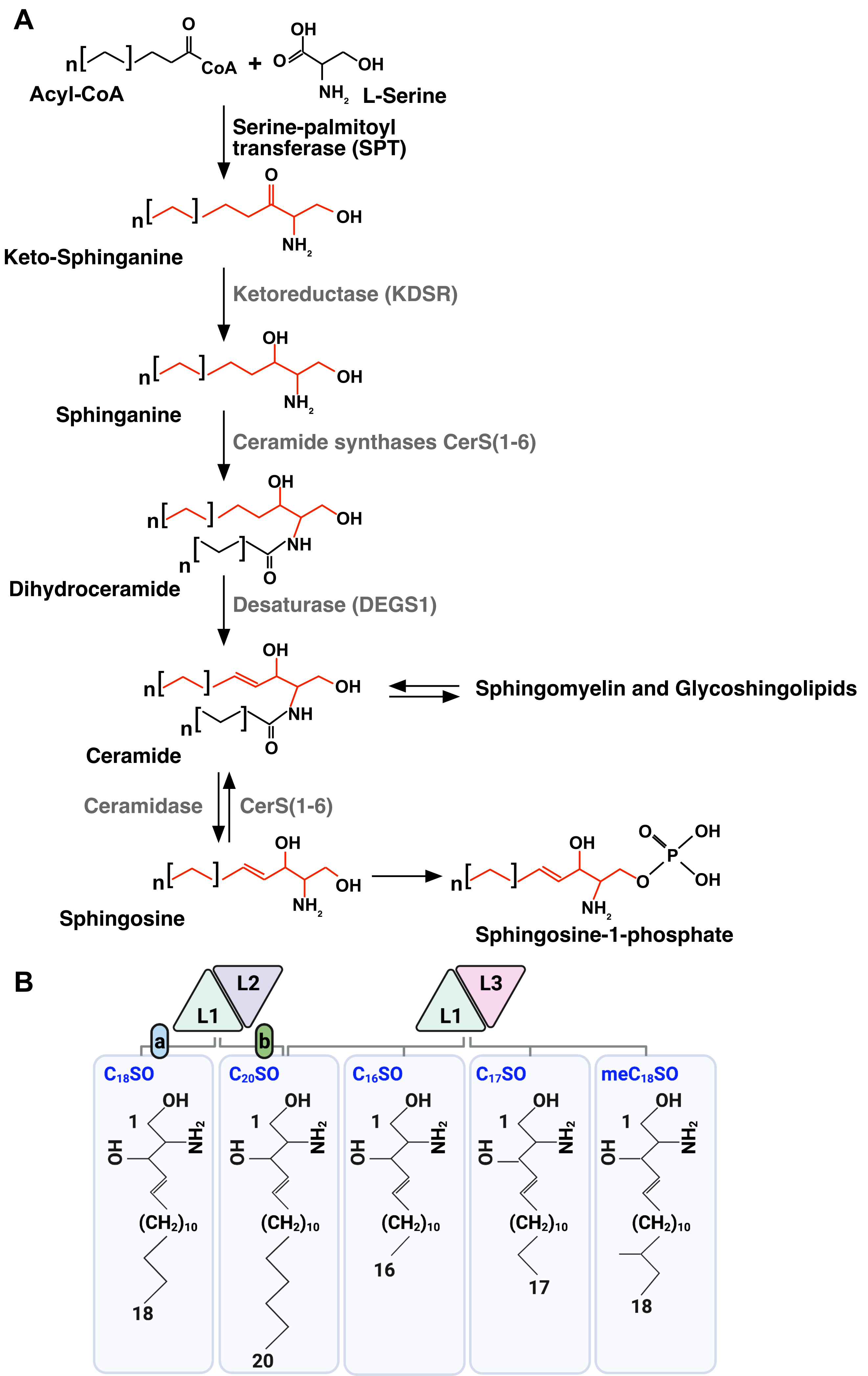
A
Acyl-CoA
CoA
L-Serine
Serine-palmitoyl transferase (SPT)
Keto-Sphinganine
Ketoreductase (KDSR)
Sphinganine
Ceramide synthases CerS(1-6)
Dihydroceramide
Desaturase (DEGS1)
Ceramide
Sphingomyelin and Glycoshingolipids
Ceramidase
CerS(1-6)
Sphingosine
Sphingosine-1-phosphate
B
L1
L2
L3
a
b
C18SO
C20SO
C16SO
C17SO
meC18SO

### Figure S1

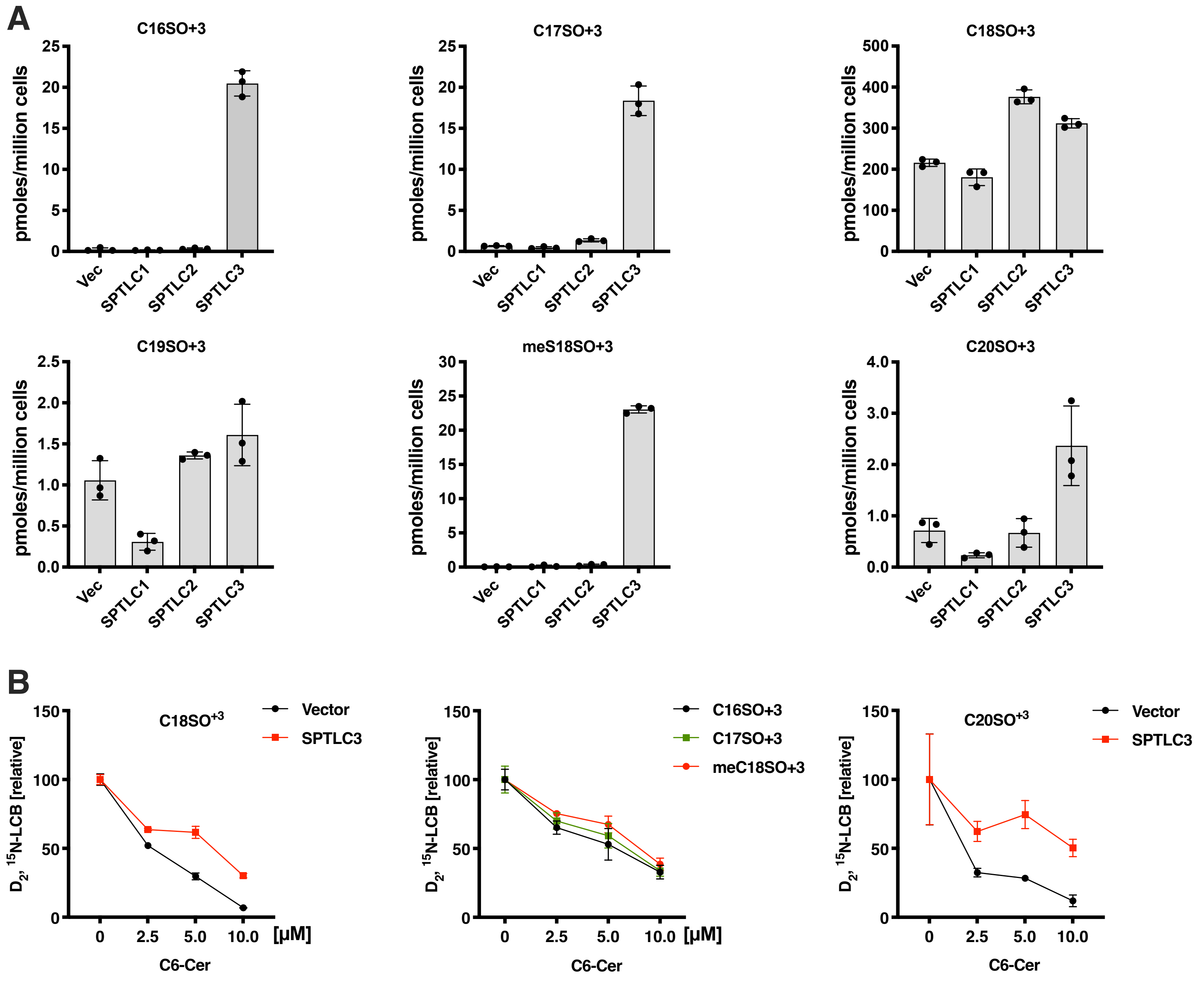

A
C16SO+3
C17SO+3
C18SO+3
C19SO+3
meS18SO+3
C20SO+3
pmoles/million cells
Vec
SPTLC1
SPTLC2
SPTLC3
B
C18SO+3
Vector
SPTLC3
C16SO+3
C17SO+3
meC18SO+3
C20SO+3
D2, 15N-LCB [relative]
C6-Cer
[µM]


### Figure S2

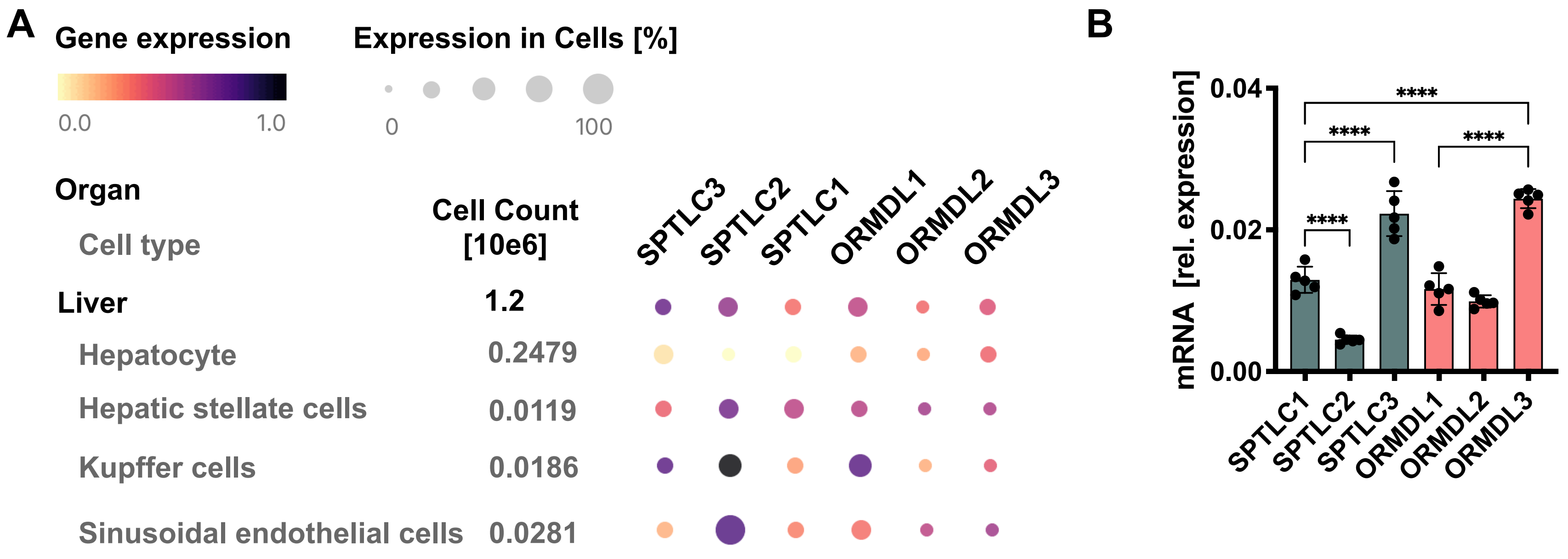
A
Gene expression
0.0
1.0
Expression in Cells [%]
0
100
Organ
Cell type
Cell Count
[10e6]
SPTLC3
SPTLC2
SPTLC1
ORMDL1
ORMDL2
ORMDL3
Liver
1.2
Hepatocyte
0.2479
Hepatic stellate cells
0.0119
Kupffer cells
0.0186
Sinusoidal endothelial cells
0.0281
B
mRNA [rel. expression]
0.04
0.02
0.00
****
****
****
****
SPTLC1
SPTLC2
SPTLC3
ORMDL1
ORMDL2
ORMDL3

### Figure S3

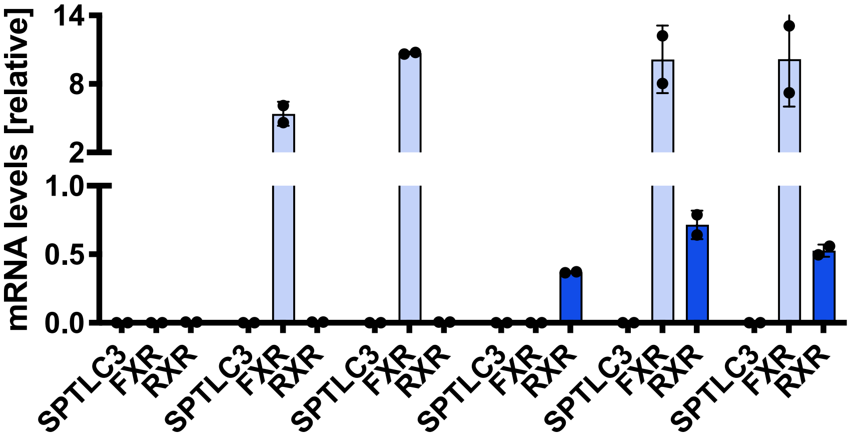
mRNA levels [relative]
14
8
2
1.0
0.5
0.0
SPTLC3
FXR
RXR
SPTLC3
FXR
RXR
SPTLC3
FXR
RXR
SPTLC3
FXR
RXR
SPTLC3
FXR
RXR
SPTLC3
FXR
RXR

### Figure S4

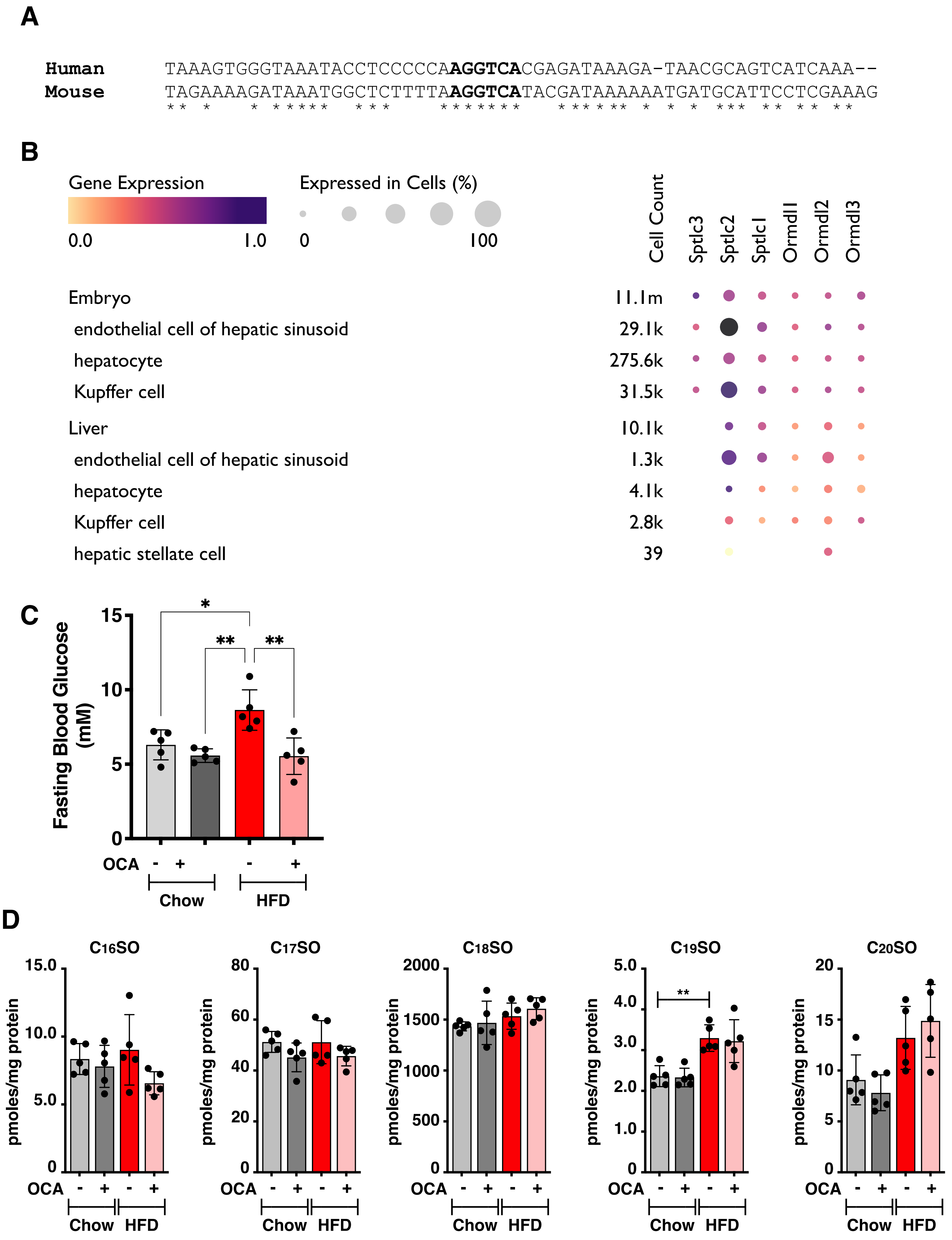
A
Human TAAAGTGGGTAAATACCTCCCCCAAGGTCACGAGATAAAGA-TAACGCAGTCATCAAA--
Mouse TAGAAAAGATAAATGGCTCTTTTAAGGTCATACGATAAAAAATGATGCATTCCTCGAAAG
B
Gene Expression
0.0
1.0
Expressed in Cells (%)
0
100
Cell Count
Sptlc3
Sptlc2
Sptlc1
Ormdl1
Ormdl2
Ormdl3
Embryo 11.1m
endothelial cell of hepatic sinusoid 29.1k
hepatocyte 275.6k
Kupffer cell 31.5k
Liver 10.1k
endothelial cell of hepatic sinusoid 1.3k
hepatocyte 4.1k
Kupffer cell 2.8k
hepatic stellate cell 39
C
Fasting Blood Glucose (mM)
OCA
Chow
HFD
D
C16SO
C17SO
C18SO
C19SO
C20SO
pmoles/mg protein
OCA
Chow
HFD

### Figure S5

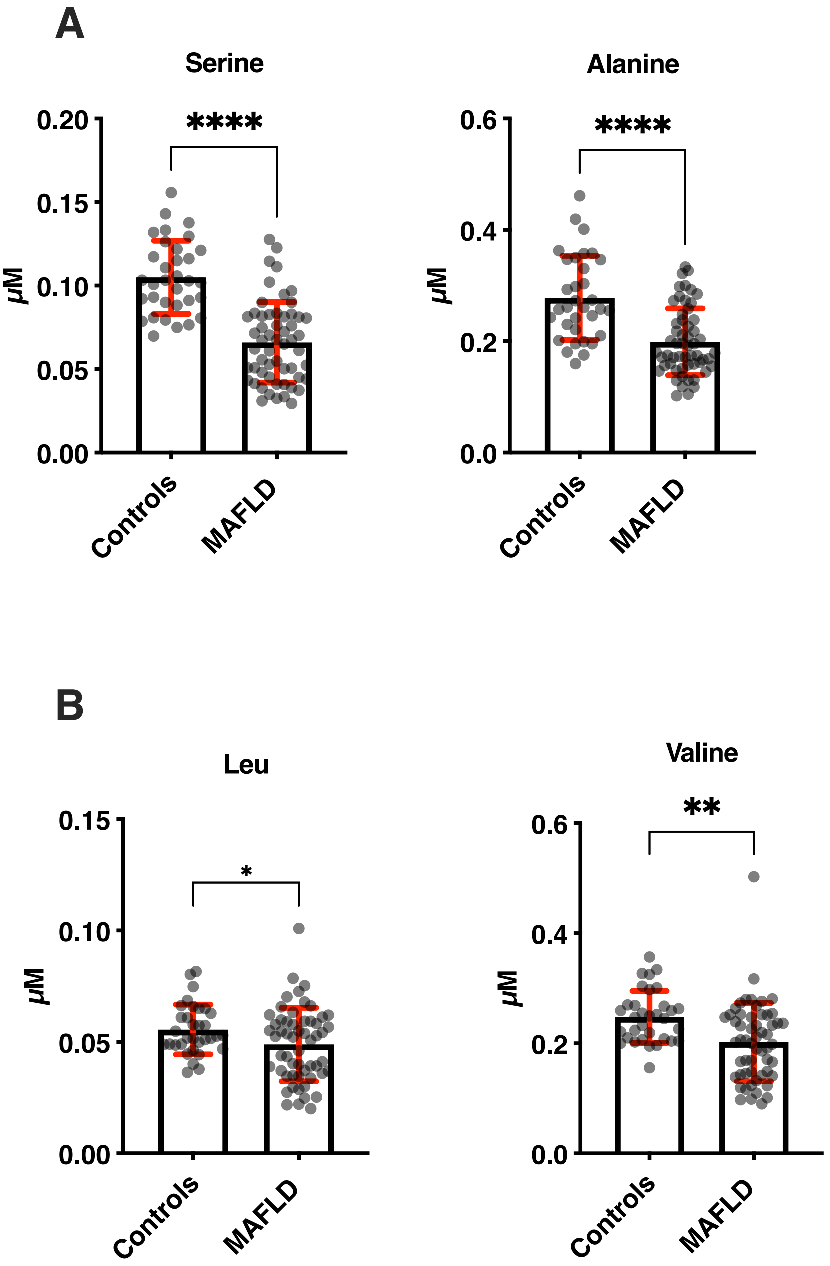
A
Serine
****
0.20
0.15
0.10
0.05
0.00
µM
Controls
MAFLD
Alanine
****
0.6
0.4
0.2
0.0
µM
Controls
MAFLD
B
Leu
*
0.15
0.10
0.05
0.00
µM
Controls
MAFLD
Valine
**
0.6
0.4
0.2
0.0
µM
Controls
MAFLD

### Figure S6
