## Supplementary Table 1 for "Dyslipidemic SPTLC3 Integrates Bile Acid-FXR Signaling with Sphingolipid Remodeling in MASLD"

Table 1 Mouse parameters

| | Cont (n=5) | HFD (n=5) | HFD+OCA (n=5) | OCA (n=5) |
|---|---|---|---|---|
| Body weight (g) | 24.20 ± 0.54 | 33.80 ± 0.86 [a] | 30.48 ± 1.48 [a] | 23.50 ± 0.57 |
| Plasma TG (mg/dL) | 121.6±2.2 | 146.7±4.6 [a] | 134.2±3.3 [a,b] | 124.5±2.9 |
| Plasma Cholesterol (mg/dL) | 60.3±4.1 | 76.5±4.0 [a] | 77.1±1.5 [a] | 64.2±4.6 |
| Liver Cholesterol ( μg/mg proteins) | 4.3±1.57 | 3.98±1.78 | 4.93 ± 2.62 | 2.06 ±0.688 |

a, p<0.05, comparison with cont; b, p<0.05, comparison with HFD
