## Supplementary material for "Dyslipidemic SPTLC3 Integrates Bile Acid-FXR Signaling with Sphingolipid Remodeling in MASLD": Suppementary Table 2

**Table 2 patient parameters**

| | Cont | MAFLD |
|---|---|---|
| Gender (F/M) | 15/17 | 22/34 |
| Age (years) | 55.03±2.62 | 55.50±2.01 |
| Plasma TAG (mmol/L) | 1.07±0.06 | 1.42±0.16* |
| Plasma Cholesterol (mmol/L) | 4.05±0.19 | 4.97±0.22** |

TAG, triglyceride. *, $p<0.05$; **, $p<0.01$; Welch's unpaired two-sample *t*-test.
