## Supplementary sequence 2 for "Dyslipidemic SPTLC3 Integrates Bile Acid-FXR Signaling with Sphingolipid Remodeling in MASLD"

ACAACCTATTCATGGGGAAATCTATGGAAATGAATGGTCTCTACGATGAGAATTTTTCCAGATTGAACTCTTCGGGAAATTTA

AGTACTTCCACGTAGACTTATTAGTAGATATTGACATGTCATTAGTCATTTGTAAAAAAATTTAATATATTGCAAATTTTTAA

ACATCAAAAAGCAAATAAAGCATTTTTATTCATTATTTGGCCAATCCAATGAAATCTCCCAAATAATAACAGATTCTGTTATT

TCTTTTCTTGGTGGTTGATGGTTCTCAAGTCAACAGCTGAAGGAGAGGTGTCATGGTGTAAGGAAAACGGCACTCACTTAGA

CCCAGACATTCTTGAGTTCATGACCCACTTCTACCACATATTATTCTGTGGCTTTGGAAAAGTCACTTAACCTGTTGAAAATTT

AGTTTTGCATGAGCATCACTCTCCATTACATGGGTAAAACTTGAGTCACTGAGAATGAACAGATTTCTGTACACTGTACAATT

CTGACAGTTCATTCTGCTCATATCCTCCCATTTTTGCCAACTCTCTATCCCCAAGTTGTTCCAAGAGCGTTTTCACAACACATT

GCCAAGCCTCTGTCCTTCCCTTTCTTGCCTATCCTCATTCTGTTATCTCCCTTTTTGGCTGCTTCATTAAGAGAGGAGCTGAGA

ATCTGGCATCTGGGCAAAACCATTACTTTGGAGAACACCACAAGGCCATTGCTAATTTTATCATCTTTGCCTTTAATGTACTA

GTGGGAAGATTTATTTCTACCTTTAATAGTAAGATATCCTAAGTAGAAGGCATGTGGGTACTGCTGAGTCTTAGTTCTGTGAA

GGTGGAGACTGTGTATGTTTTATTCCTTGTGAGAACCAAGGACTCAGCAAATACAGAGTACTTATTCAAACATTTTCTGAGGT

TACTAGTTGATCCAAAATGTATTTAGTTGTTCAGAGATGCCAAATCTTGCTCAATATTTAGTCTGTGTTTTAGTATTGCTCCAA

***cis*-element 1**

ACCTTAAACCTAAACCTACCACTGACTTCAACAGAACTCTTCGCTTATTTTAACTGGAGATGGTATGAAGCAATGAGTCACC

ACTGACTATACATACCAGCAGTGCCTGATGCCAAATGGTTGGTGTGTTAATATAATTTTGTACACACAACAATGCACACATTC

CAATACGGTAAGTAAAATAGAAAAGTTTCAAGAGTGAGAGCACGTAGTAGATATTTCTCAGAAATCTATGTGTGATTACCTA

ACCTTAGCAATTGCAATTAGGGGAAGGAAACGGAGAAAGCAGAATTCAGCTGTGTTTTAAGATTTATTTTATATATACCTTTT

GTCCCTTGTAAAAAGAAAATTTGAAAAAGAAGAAGAAAAAGAAGAGGAGGGTGGGGGAGGAAAGGGAAAAGGACAAATG

TGAAAAGAGGAAAGAGATGGAGAAGAAGAAAAGAAGAAAGAAAAGAGACATCTAATGTTAAAAAAGTTTTATTTCAAAAT

CTTTATTAAAAAAATGCTTTGCACTGAGCAATAGAGCTCAGGGAAAATAACGCCCTCTCTTCAACTTGTCTTTCTAAGTAACG

TGAGGCAGTTGGGCCAACAATAGTGCTCCCCTCTACCTCTTCATCTCTCCATTTCCTAAATCTTAGAAGAAAATAAACCTACA

***Pr.SPTLC3-GLuc***

AGAGTTTAGCCCCCAAATCTCAGTGAAACAGCCATGGAAGGGAGAGACCAGTGAAAATCTACTTCTACTTTGATGATAAAGG

TTGAGAGAAATGCGATAAAATTTTTGAACTTGAGACACAAACTGGGCTGAACACACTTGGTATTAACTCCGTCTTTCACCTTG

GCCTCAGGCCTTATCTTATTCCTCCAGCATTCCTTGGGATGGTAAAGGCAGGTACCCTCATATGCAAGTCATAGTTAGGAGAA

GCGTTAGCATGTCCACCTCCAAAAGCTACATTAATCTGGAGTAGAACTGAACCCTGTCCAAGGGAGCTGAGTTACAGGACAT

GAGGCTCTGGAAACAGGGGCCATTTCTTTCTTTTTGTTTGTGTGTGTGTGTTTGATTGTTTGTTTTTGTCTACTGCCCCAATAA

ATTGGTTTCTTTTCTTAGAGTCTGATGCTAAAGAAGTCTCGACTCAGGAAAACCACAAGGGTGGACTTGGGAAGACTCAACA

GGTTGTAATGACTCTCCAGCAAATCATGTCACCATGCCATTTGATGATGGTAGGAAGAAACAAAGCCTTGAAGATTCATGCA

TTGAGAGAATAAGAGTTAGTGAGTAAACCAGATGTGTTTCAGGACTGGGACCCTGCAGTCATCAACCACCTCACACCTGCGT

CAGTAACAAACCCAAGCTACAGCATGATTTTTGTTCTACCTCAGCAGATATTTTATAGATGGAAAAACTGAGGCATAAAGTG

***cis*-element 2**

GGTAAATACCTCCCCCA<u>AGGTCA</u>CGAGATAAAGATAACGCAGTCATCAAATATTCTGTCTCCTCAACATCTGTCTTACTGGA

ATTCTTGATGGATTGGTACCCCCTTATTATAGTACTGATTTTAACATAGGTTGGTTTTGAATACTGGGTTTTTAAAAATCTTTT

CAAATCATTTGTAAATTATGGAAATGTATACATCACATGAAACTTACCATTTTAATCATTTTTAAGTGCACAGTTTAGTTTAA

GTACATTCACATTGTTGTGCACTGATCCATCACTATACCTTTTTCATCTGCTCCAGCTCAAACTCTGTACCCATTAAACAATAA

CTTCCCTTTGCCTCATCCCTCCAGGCCCTGGCAACCACCTCTGAATGCTATTTTATAGCAAAAAGAAATATTAACTTTTGAGCT

TGGTTTACAACCAAAAACCACAATTACATGCAGGAGAGAGGGGGGTGGGAAAAAGAGGTTAACTCAGAACTCAGTTACTTAG

AAGACTCTGTGAGTGTGCTGTGTGTGTGTGTCTTTAAAAGACTCTCCACCTCCCAGCCCGCCTCCTCACACTTTGCCACTGGG

**TSS**

TTGTTCAGTCCCCAGGTTCCCTCAGTCCCC<u>A</u>GAAGGAGCCAGCATGGACAATCTCCTTTACAGTTTCGGAAGCAGGTTTGTTG

CCATGGAGTTCACATTTTGACGGGAGTTGAGAAGTATAAAGGTAACCATTTGTTTTAGTTTCAACGATCTGACAAAAAGATA

GGCTGTTGCTCTTCTTCTGGAAAAGCCTGATTGGTAAGATTCCTTTAAGGGCTCAGCCCCAAAGAGCTTTATCCCATCCCCTC

**SPTLC3 Start**

GCAGACTGAAAACTAAAGCCTGCAGAGACCTCTGAAGGAAAACCTGTCCCGGGCTCTGTCACTTCACACCC<u>ATGGCTAACC</u>

<u>CTGGAGGTGGTGCTGTTTGCAACGGGAAACT</u>TCACAATCACAAGAAACAGAGCAATGGCTCACAAAGCAGAAACTGCAC

AAAGAATGGAATAGTGAAGGAAGCCCAGGTAAGAGGCACTCTCCCCTACTCTTCTCTGAATTACCTGAACGTTTCAATATTT

CTCTGTGAAGATATTTGAGGCTGTCCTAACTTAGAGTTCATCTTATGAAGGGTTTTTACATGTTTTGACTGTT
